# The reproduction of Gram-negative protoplasts and the influence of environmental conditions on this process

**DOI:** 10.1101/2021.11.25.470037

**Authors:** Dheeraj Kanaparthi, Marko Lampe, Jan-Hagen Krohn, Baoli Zhu, Andreas Klingl, Tillmann Lueders

## Abstract

Bacterial protoplasts are known to reproduce independently of canonical molecular biological processes. Their reproduction is shown to be mediated entirely by the physicochemical properties of cell constituents. However, the physiochemical properties of the cell constituents are influenced by the environmental conditions like salinity, salt composition, and mechanical stresses experienced by a cell in natural environments. The influence of such environmental conditions on protoplast reproduction is seldom investigated. Here, we studied protoplast reproduction in their native environmental conditions. Contrary to the previous perceptions of protoplasts reproducing in an erratic manner, cells in our study reproduced in a defined sequence of steps. The process of their reproduction can be explained by an interplay between intracellular metabolism, the physicochemical properties of cell constituents, and the nature of cations in the growth media. We observed a minimal leakage of intracellular constituents during protoplast reproduction, suggesting an efficient reproduction. However, the efficiency of reproduction is determined by the environmental conditions. Under favorable environmental conditions, protoplasts reproduce with nearly similar efficiency to cells that possess a cell wall. In short, here we demonstrate the simplest method of cellular reproduction and the influence of environmental conditions on this process.

## Introduction

Bacterial species typically possess a peptidoglycan cell wall, which imparts structural rigidity and forms the foundation of their morphology^1^. In addition to its structural functions, the cell wall plays a vital role in maintaining proper turgor pressure, transporting nutrients into the cell, and facilitating reproduction ^2^. However, bacteria are known to lose their cell wall and transform into their protoplast state when exposed to cell wall-degrading enzymes and bactericidal antibiotics ^3,4^. Although this transformation is thought to make cells susceptible to changes in environmental conditions like osmolarity ^5^, it is also known to confer some advantages to the bacteria ^3,6,7^. Most of the antibody-binding sites are located on the surface of the cell wall ^8,9^. When these cells transition to their protoplast state, they avoid being detected by the immune system of the host cell, facilitating the unchecked proliferation of pathogens within the host ^10^. Given these clinical implications, most research on protoplasts primarily focuses on understanding either the clinical implications of this transformation ^11–13^ or the intracellular processes within the protoplasts ^14,15^.

In contrast to our conventional understanding, some recent studies showed that the transformation of bacteria into their protoplast state could happen in natural environments without antibiotic stress ^6,16^. This transformation is observed to have been induced by environmental conditions like osmolarity and exposure to lytic phases ^6,16^. This transformation of cells into the protoplast state is shown to confer cells with phase resistance, increased survival rates in natural environments, and receptivity to incorporating DNA from the surrounding environment ^6,7,17^. Apart from the well-documented clinical implications of bacterial transformation into protoplasts, these studies imply the ecological benefits of such transformation ^6,16,17^. However, we currently lack a good understanding of the ecological aspects of this cell state, especially how the environmental conditions influence the protoplast life cycle.

Protoplasts are known to reproduce independent of canonical molecular biological processes^14^. Reproduction in such cells is shown to have been primarily mediated by the physiochemical properties of the cell constituents, like the membrane fluidity and ratio of cell volume to surface area ^15,18–20^. The physicochemical properties of cell membranes, like the fluidity and spontaneous curvature, are influenced by external conditions like osmolarity, the nature of the cations, and the mechanical strain experienced by a cell in natural environments ^21–24^. This suggests that conditions surrounding the cell exert a considerable influence on the morphology and reproduction of protoplasts. Despite this possibility, much of our current understanding of the protoplasts is a result of studying them under well-controlled environmental conditions rather than in their native environment ^4,14^. In this study, we examined protoplast reproduction in their native environment and how various environmental conditions influence this process.

The cytoplasmic osmolarity of a non-halophilic bacteria ranges between 300-500 mosM ^25,26^. This was a result of the densely packed intracellular environment with proteins and nucleic acids. Maintaining this high intracellular osmolarity is necessary for the cells to absorb water and nutrients from the surroundings ^27^. One essential function of the rigid peptidoglycan cell wall is to counteract the uncontrolled expansion of cytoplasm due to its high osmolarity ^28^.

Given that the protoplasts lack a cell wall, they are prone to osmotic lysis ^5^. Hence it is necessary to grow them in an isotonic media ^14,18^, and their distribution in natural environments seems restricted to mildly saline environments ^11–13,17^. To emulate these conditions, we grew the bacterial protoplasts in media containing 7%Dead Sea salt (DSS) at 30°C. DSS was used in our incubations rather than pure NaCl to replicate the complex salt composition of the natural environments. In the below sections of the manuscript, we present the lifecycle of protoplasts under environmental conditions close to their native habitat.

## Results

### Generation and physiological characterization of protoplasts

We transformed *R. sphaeroides (RS)* into its protoplast (*R. sphaeroides-protoplast, RS-P*) state by the procedure described in the methods section. This transformation was evident from the change in the morphology of cells from coccoid to relatively bigger spherical cells (Figure 1). The absence of the cell wall was also confirmed by transmission electron microscopy (TEM) (presented below). *RS-P* did not transform back into its native state when the use of lysozyme and penicillin was discontinued. But a step-by-step reduction of salt content in the media or gradually replacing salt with an equivalent concentration of sucrose led to the transformation of cells into their native state with a cell wall. No reversal of *RS-P* to native-*RS* state was observed in control incubations where the salt concentration is consistently maintained at 7% or in sugar-rich media with lysozyme and penicillin.

**Figure 1.**
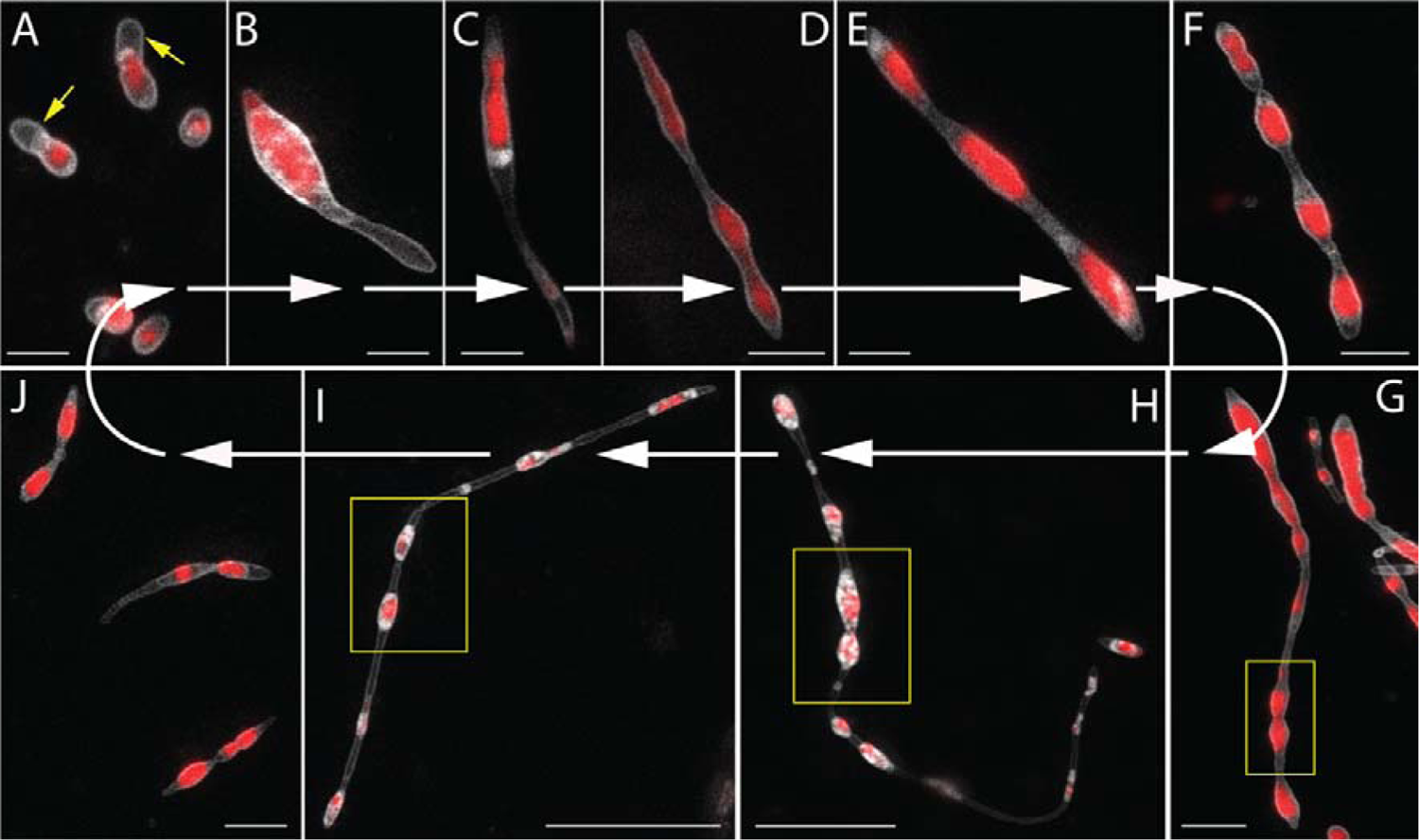
The lifecycle of *RS-P* All panels show STED microscopy images of cells stained with FM^TM^5-95 (membrane, white) and PicoGreen^TM^ (DNA, red). White arrows guide the eye in following the life cycle progressing from A to J. Arrows (yellow) in A highlight a cell with outer membrane extensions (are also seen in B & C). The steps involved in forming outer membrane extensions are shown in Figure 2. Boxed regions in G – I highlight cytoplasmic compartments within the filamentous cell undergoing binary fission (in sequence from G-I). J: Individual daughter cells with membrane overhangs formed from the fragmentation of filamentous cells. Also, see Figure S2 for phase-contrast images. Scale bars: 2μm (A-G & J) and 10μm (H & I).

Substantial differences in the physiological behavior were observed between *RS-P* and “wildtype” *R. sphaeroides*. *RS-P* exhibited similar superoxide dismutase (SOD) activity under aerobic (14.8 ± 1.3 U/ml) and anaerobic (15.2 ± 2.1 U/ml) conditions. In contrast, the SOD activity of WT *R. sphaeroides* depended on environmental conditions. Considerably lower cytoplasmic SOD activity was observed under aerobic growth conditions (8 ± 0.04 U/ml) compared to *RS-L*. Negligible to no SOD activity was observed when *R. sphaeroides* was grown in anaerobic media (0.16 ± 0.05 U/ml).

Fermentation products like lactate, butyrate, and acetate were observed when *RS-P* was grown in well-aerated media containing glucose (Figure S1). Such volatile fatty acid accumulation was not observed in *R. sphaeroides* incubations under aerobic conditions (Figure S1). All the experiments were replicated (at least n=5), as indicated in the individual methods sections. The lifecycle and other data presented below were repeated numerous times over the course of this work.

### The life cycle of *RS-P* in 7%DSS-TSB

Once we obtained stable protoplasts, we transferred them into fresh growth media containing 7% Dead Sea salt (DSS). We then monitored these cells at regular intervals using various microscopic techniques. The lifecycle of these cells is shown in Figure 1.

During the lag phase, growth was restricted to the outer membrane of the cell, which expanded to form a hollow filamentous extension (Figure 1A-C, 2 & S2). The length of these extensions varied from cell to cell, ranging between 0.5 to 5 μm (Figures 1A-C, 2 & S2). Most cells in this growth stage were observed to have developed hollow membrane vesicles within their cytoplasm and in the periplasmic space between the outer and inner membranes (Figure 2). These vesicles initially appeared as tiny buds attached to the inner cell membrane (Figure 2D, arrow), followed by their expansion into or out of the cytoplasm (Figure 2D-F).

**Figure 2.**
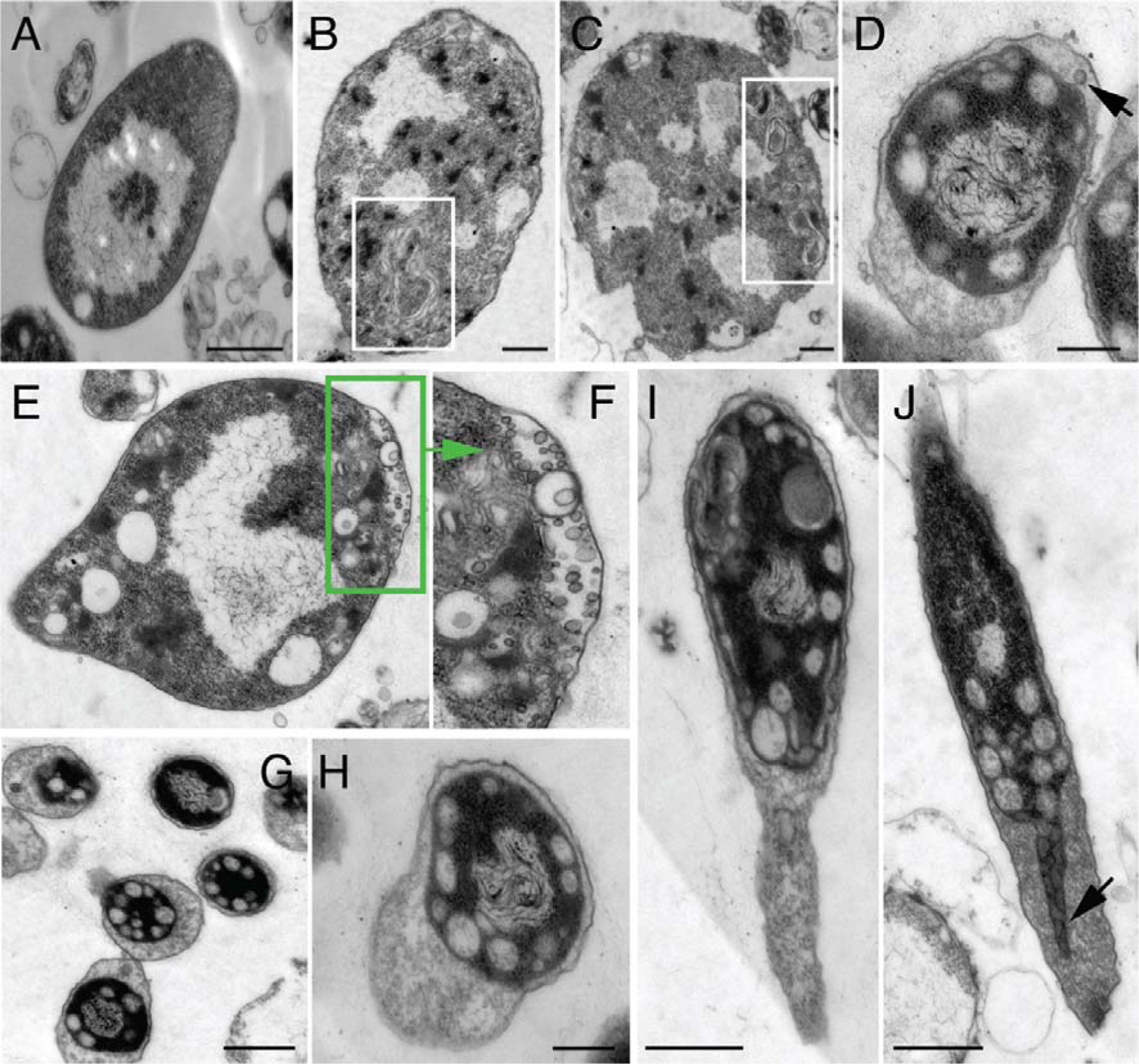
TEM images of early growth stage *RS-P* cells. Images A – C show cells with folded excess membrane within the cell (highlighted regions). Image D shows *RS-P* cells with intracellular vesicles. The arrow in D points to the inner membrane undergoing budding into the periplasmic space. A cell with hollow vesicles in the periplasmic space was shown in E. Image F shows the magnified periplasmic region of the cell in E. Images G – I show cells with the elongated outer membrane. Image J shows the cytoplasmic compartment’s growth, filling in the void left by the outer membrane (arrow). Scale bars: 0.5μm (A-J).

During the early-log phase, with an increase in the growth rate (Figure 3), the volume of the cytoplasm increased, gradually filling the hollow template created by the outer membrane (Figure 1C&D, 2J, and S1A-C). This led to the transformation of cells in sequence from spherical to tubular (Figures 1, 3, and S2), then to dumbbell-shaped cells with cytoplasm mostly restricted to either end of the cell (Figure 3, early log-phase). In the subsequent growth stages, cells developed surface depressions along their length resembling tubular vesicles subjected to curvature-induced destabilization at their surface (resembling Rayleigh-Plateau instability) (Figures 1D-G, 3A, & S3). These surface depressions were formed due to the cytoplasmic compartment surrounded by the inner membrane undergoing binary fission (Figure 1D-H & G-I).

**Figure 3.**
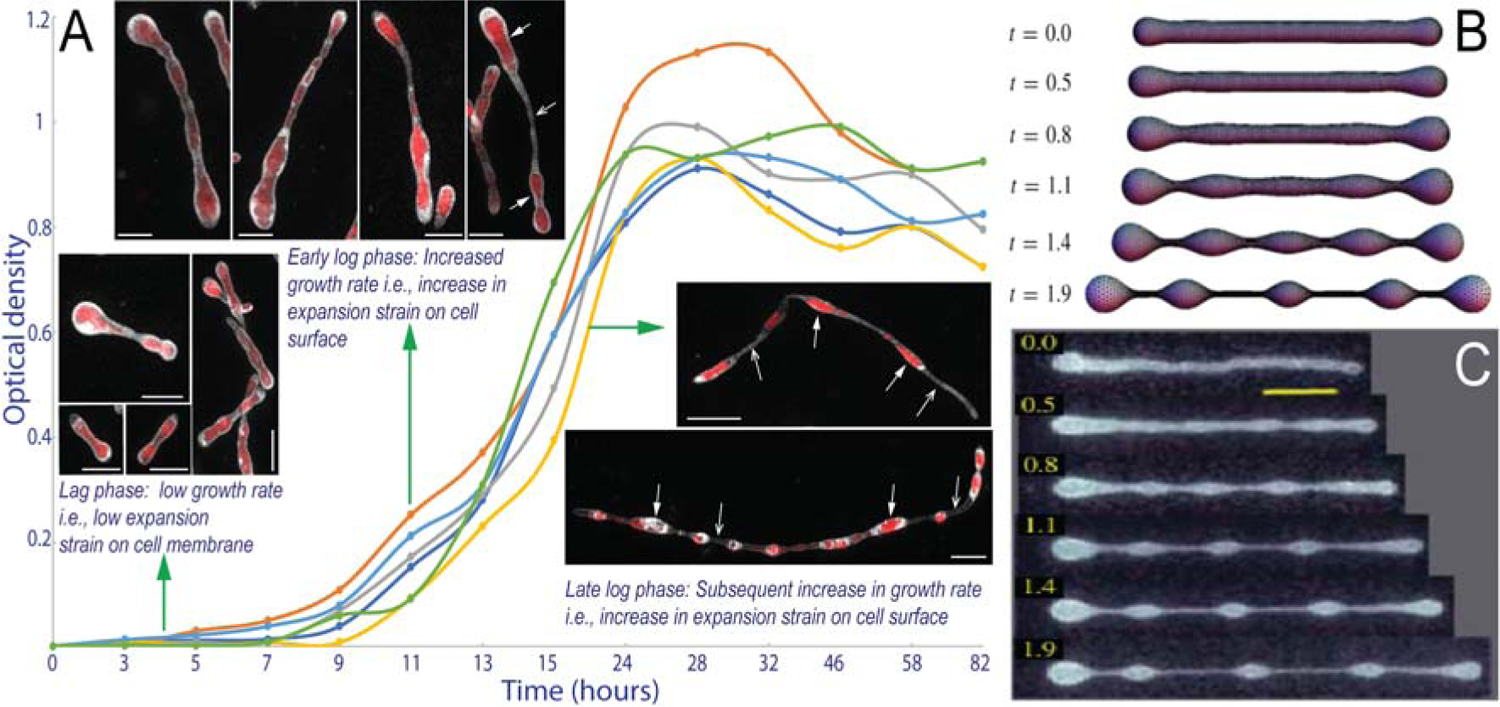
*RS-P* cells in different growth stages and comparison of their morphology with LVs. Image A shows the growth curve and morphologies of *RS-P* in different growth stages (individual plots represent multiple repetitions, n=6). Cells were stained, imaged, and color-coded, as in Figure 1. Closed and open arrows point to cytoplasmic compartments and hollow outer membrane connecting the cytoplasmic compartments, respectively (Figure 4). Images B and C show theoretically predicted and experimentally observed morphologies of vesicles subjected to elongational stress over time (t) (images B & C were originally published by Narasimhan et al., 2015 ^29^, reproduced here with permission from Cambridge University Press). A similar comparison of *RS-P* cells with LVs undergoing expansion is shown in Figure S3. Scale bar: 1μm (lag & early log-phase), 10μm (late log-phase).

Over time, these surface depressions grew deeper, and two adjacent cytoplasmic compartments were linked to each other only by a hollow strand of the outer membrane (boxed regions in Figures 1H&I and 3 – late log phase). The rupture of the outer membrane connections led to the formation of individual daughter cells (Figure 1J). This sequential transformation of cells was not observed when we repeatedly transferred log-phase cells into fresh media every few hours to keep them in a prolonged log phase. Cells in these experiments grew into longer and thicker filamentous cells without developing surface deformations (Figure S5) or transforming into a “string-of-beads” morphology. The cytoplasm of these cells was observed to have multiple intracellular vesicles (Figure S5C-E). In the stationary growth phase (5-8 days from the start of the incubation), cell morphologies were a mix of all the above-described morphologies and a considerable amount of membrane debris (Figure S5).

### Membrane phase separation in *RS-P*

Staining *RS-P* cells with both universal membrane dyes (FM^TM^5-95) and a dye specific for the liquid-disordered membrane (L_d_) (*FAST*^TM^DiI) showed the existence of two distinct phases within the cell membrane: liquid-ordered (L_o,_ red) and liquid-disordered (L_d,_ green) phases (Figures 4, and S7). During early growth stage, the cell membrane underwent phase separation, and both L_d_ and L_o_-membrane underwent invagination to form hollow buds. All the buds extending out of the cytoplasm into the periplasmic space were exclusively composed of the L_o_-membrane (Figure 4F, arrow), and buds extending into the cytoplasm were composed of the L_d_-membrane (Figure 4A-E). Over time we observed a gradual enrichment of a more fluid L_d_-phase within the inner cell membrane and a more rigid L_o_-phase in the outer cell membrane (Figures 4 and S7).

**Figure 4.**
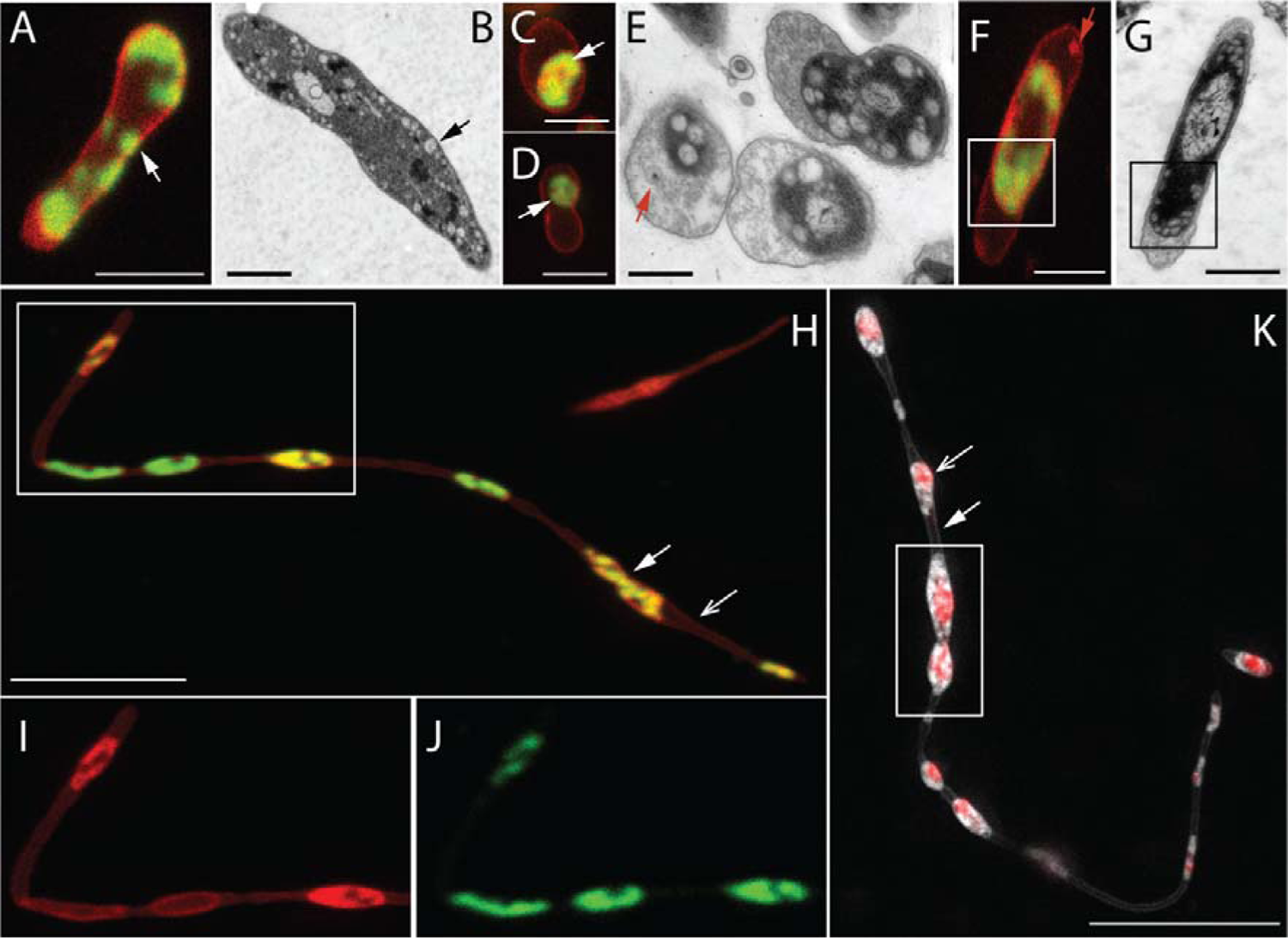
Membrane phase separation in *RS-P*. Cells in images A, C, D, F, H, I, & J are stained with FM^TM^5-95 (all membrane, red) and *FAST*^TM^ DiI (L_d_-membrane, green). A & B: confocal and TEM images of early growth stage cells. Arrows in image A point to regions of the cell with L_d_-membrane. Arrows in image B point to similar intracellular vesicles. C - G: confocal and TEM images of later growth stage cells. Arrows in C & B point to the cytoplasmic compartment, exclusively enclosed in the L_d_-membrane. The Arrow (red) in E point to the vesicles in periplasmic space. The arrow (red) in image F shows similar periplasmic vesicles composed of L_o_-membrane. H: Late log-phase filamentous *RS-P* cells (see Figure S7 for multiple optical sections of this stack). I & J: Magnified regions of the boxed region in H in individual L_o_ & L_d_-membrane channels. K: *RS-P* cell similar in morphology to the cell shown in I. The cell in this image is stained with membrane and DNA stain, as in Figure 1. A comparison of cells in H & K shows the inner cell membrane enriched in L_d_-membrane. Scale bars: 1μm (A, C, D, & F), 500nm (B, E & G), and 10μm (H & K).

### Influence of environmental conditions on *RS-P*’s morphology

To understand the influence of individual salts on the morphology of *RS-P*, we repeated the above experiments replacing DSS in the growth media with an equivalent concentration of either only divalent or only monovalent salts like MgCl_2_ or KCl/NaCl. Cells grown under these conditions exhibited morphologies similar to those grown in DSS (Figure 5). However, considerable differences were observed in their size – considerably thicker and longer filamentous cells (Figure 5A-D) and considerably smaller individual cells (Figure 5E-H) were observed when cells were grown in media containing 7%KCl and 7%MgCl_2_, respectively. Thick tubular or long filamentous cells with diameter >1-2μm were rarely observed in incubations with MgCl_2_, often only transiently during the log phase (Figure 5F&G). When grown on an orbital shaker, hardly any filamentous cells were observed in our incubations (Figure 5I-L). Instead, most cells during all growth stages are unicellular or cells undergoing binary fission. Filamentous cells were rarely observed (Figure 5I-L). Treatment of *RS-P* with 5mM of 3-methoxybenzamide ^30^ did not result in any apparent morphological changes compared to the control incubation (Figure S8).

**Figure 5.**
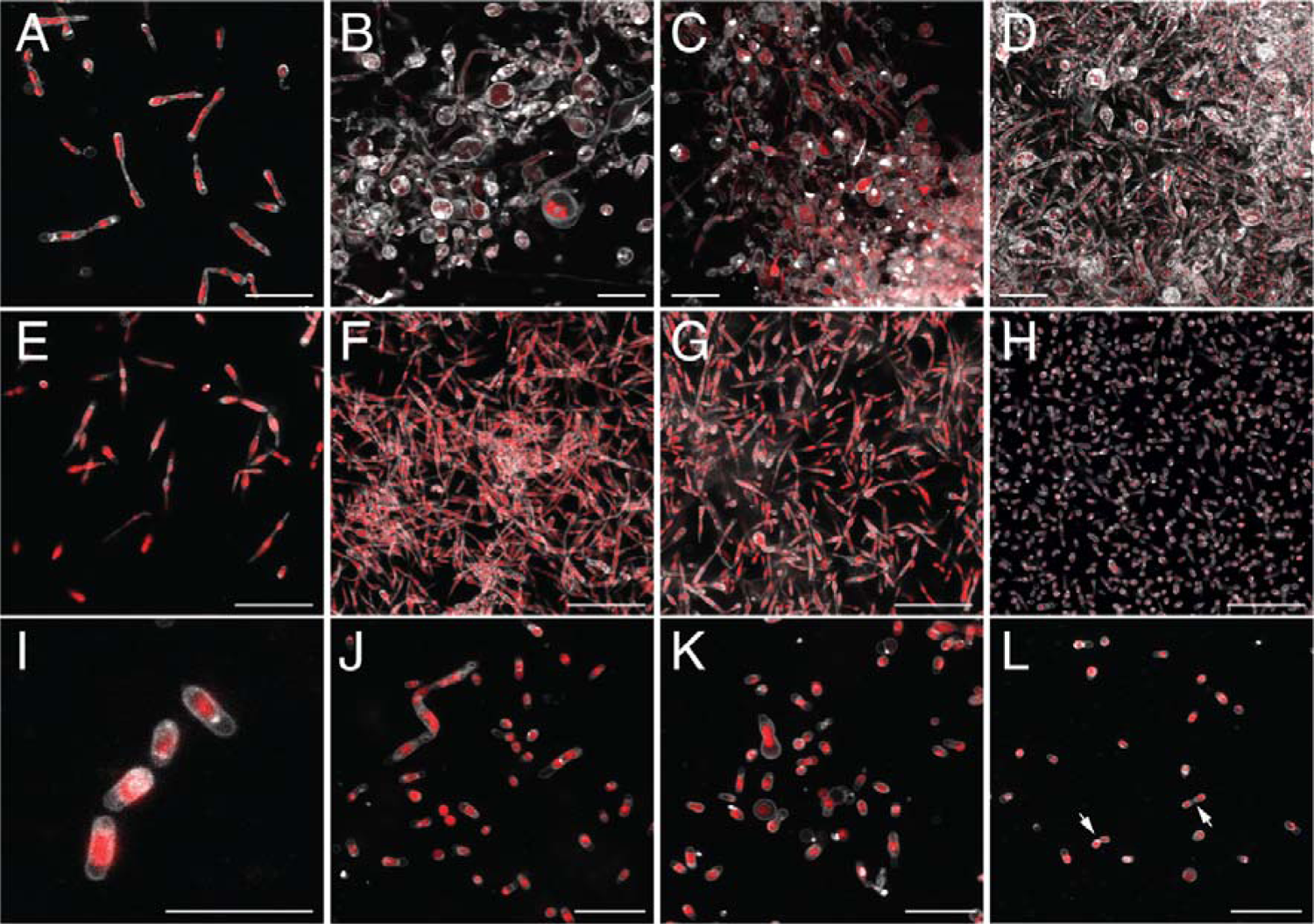
Influence of salt composition on *RS-P*’s morphology. Images A – D shows *RS-P* cells grown in 7% KCl-NB from early to late growth stages. Images E – H show *RS-P* cells grown in 7% MgCl_2_-NB from early to late growth stages. Images I – J show *RS-P* cells grown in 7% DSS-NB on an orbital shaker. Arrows in image L point to cells undergoing binary fission. Cells were stained, imaged, and color-coded, as in Figure 1. Scale bars: 10μm.

### Reproductive efficiency and viability of *RS-P* daughter cells

The concentration of extracellular DNA during the stationary growth phase was used as a proxy to determine the leakage of intracellular constituents during reproduction. The results of this quantification are presented in Figure 6. The concentration of extracellular DNA in all our incubations was an order of magnitude lower than the intracellular DNA. Quantifiable but no significant differences in the concentration of extracellular DNA was observed between *RS-P* and the native *R. spheroides* cells, which possess a well-functioning molecular biological regulatory process (Figure 6A). Neither were significant differences observed between cells grown in media containing different salts. The reproductive efficiency was marginally higher when *RS-P* was incubated on an orbital shaker rather than under static conditions (Figure 6B). Staining and imaging of the stationary growth phase incubations also suggest the absence of extracellular DNA (Figures 5, 6C, and S6).

**Figure 6.**
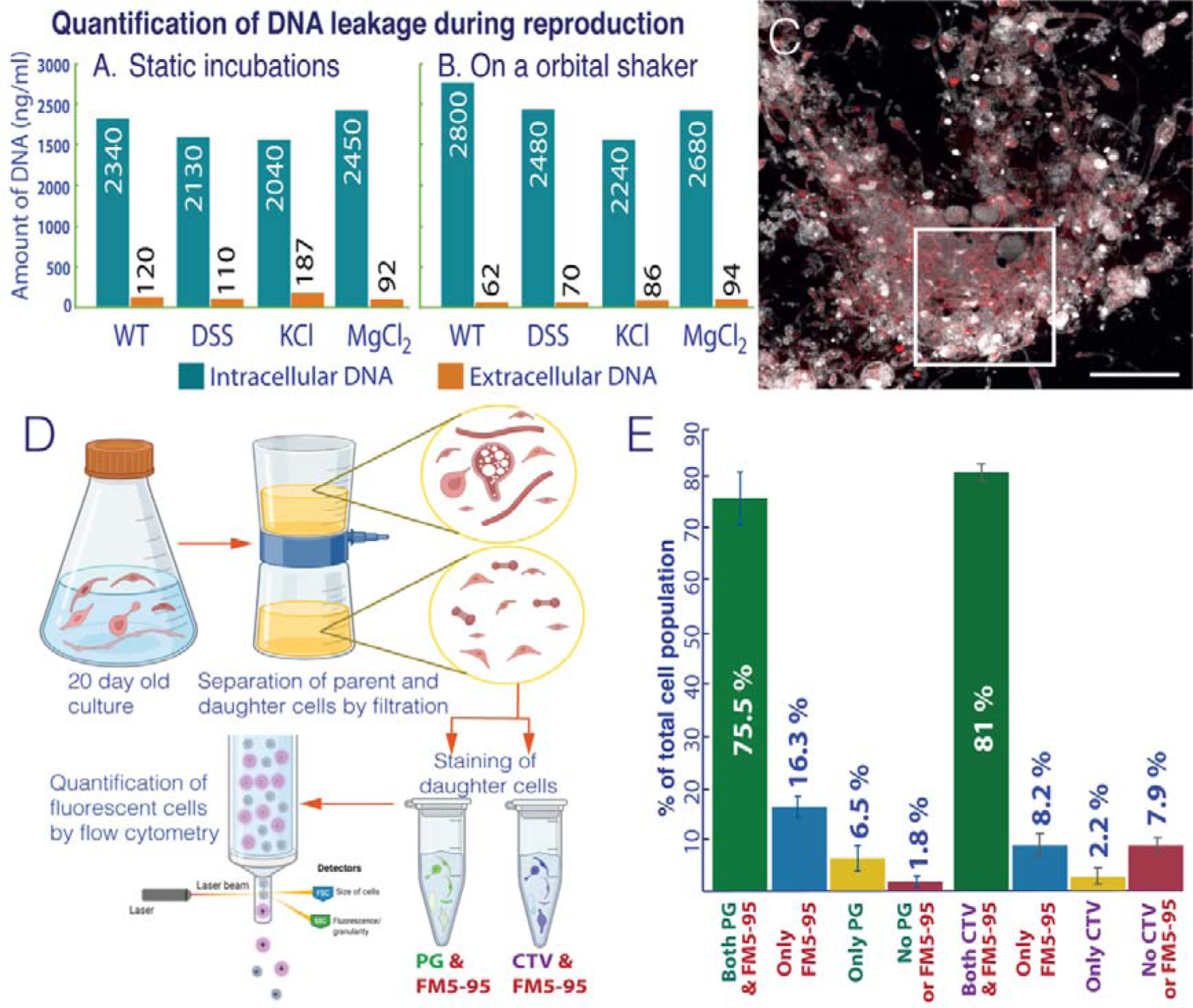
Reproductive efficiency and viability of *RS-P* daughter cells. Graphs A & B show intracellular (blue bars) and extracellular (orange bars) DNA concentrations (n=5 for each condition). A: *RS-P* grown under static conditions. B: *RS-P* grew on an orbital shaker. Annotations: WT (wild type, *R. sphaeroides* with a cell wall grown in NB); DSS, KCl & MgCl_2_ – *RS-P* grown in NB with respective salts. Image C shows *RS-P* cells grown in 7%DSS-NB in the late growth stage (Scale bars: 10 μm). Cells in this image are stained, as described in Figure 1. D: Schematic workflow of the experiment to quantify live daughter cells. Cells were stained with different combinations of FM^TM^5-95 (cell membrane), PicoGreen^TM^ (PG, DNA), and CellTrace^TM^ Violet (CTV, cell integrity) (n=5, in the figure but multiple repetitions over the course of the work). E: The percentage of *RS-P* daughter cells identified positive for different viability markers in flow cytometry. The original flow cytometry plots are shown in Figure S9.

The metabolic viability of *RS-P* daughter cells was determined by staining daughter cells from the stationary growth phase with dyes specific for cell membrane, DNA, and cytoplasmic activity (Figure 6D). Quantification of live cells (cells with intact membrane, intracellular DNA, and cytoplasmic activity) was done by flow cytometry. Our results suggest that most of the daughter cells (< 70%) received DNA from the parent cell and are physiologically active (< 80%) with intracellular enzyme activity (Figure 6E). Subsequent transfer of daughter cells of different ages – 10, 20, or 100-day-old cultures either directly or after passing through a 0.45μm filter (to separate daughter cells from larger parent cells and membrane debris) into a new media always resulted in growth. Transfer of daughter cells into minimal salt media^31^ containing glucose as a sole carbon and energy source resulted in growth (Figure S1).

## Discussion

The above-described life cycle of *RS-P* could be explained by the lack of molecular biological coordination between physiological processes within the cell, the biophysical properties of its cell constituents, growth rate, and environmental conditions. This loss of intracellular coordination between cellular processes is evident from the excessive lipid synthesis, production of volatile fatty acids from under aerobic conditions, and high intracellular SOD while fermenting glucose. In a normal functioning cell, the rate of lipid synthesis is tightly regulated by its growth rate ^32^. In contrast, *RS-P* was in a perpetual state of excess lipids. This is evident by the excessive membrane within *RS-P* during all growth stages (Figures 1, 2B, 2C, S2A, and S4).

The flow of electrons in the electron transport chain is known to produce reactive oxygen species (ROS) ^33^. Cells were known to synthesize catalase and SOD enzymes to prevent ROS-related oxidative damage. Such enzymes are selectively expressed only during the availability of terminal electron acceptors and were repressed during growth by the fermentation ^33^. High and low cytoplasmic SOD activity of *RS,* when grown under aerobic and fermentation conditions, aligns with our current understanding of SOD-related gene expression. In contrast, *RS-P* exhibited high cytoplasmic SOD activity irrespective of its physiology or the availability of terminal electron acceptors. Moreover, we observed the accumulation of volatile fatty acids like lactate, butyrate, and acetate in growth media when *RS-P* was grown under well-aerated conditions (Figure S1). Approximately 30-35% of the glucose was converted to fermentation products. This transformation of glucose to volatile fatty acids happens only under anaerobic conditions when the growth media lacks terminal electron acceptors, and the formation of such compounds was not expected during aerobic growth. No fermentation products were observed when *RS* was cultured under aerobic conditions (Figure S1). These differences in the physiological behavior between *RS-P* and *RS* also suggest the loss of mechanisms to sense the environmental conditions or the loss of molecular biological coordination required to adapt cellular physiology to the ecological conditions. A loss of molecular biological coordination leading to chaotic behavior of cells ^4,14,20^, like the excessive lipids synthesis ^34^, and lack of ability to assemble a peptidoglycan cell wall, despite upregulation of the genes for synthesis of a cell wall components ^35^, was previously reported in protoplasts. Although our observations suggest a lack of coordination within the cytoplasmic processes and are in tune with the previous studies ^4,34,35^, further substantiating this claim requires an in-depth molecular biological investigation, which is beyond the scope of understanding the protoplast reproduction in natural environments.

Morphological features observed in *RS-P,* like the outer membrane extensions, hollow vesicles within the cells, and transformation of tubular cells to a string-of-beads morphology (Figure 3), can be explained as a mechanism of accommodating this excess membrane. Transfer of lipids synthesized within the cytoplasm to the outer membrane and its subsequent expansion into a hollow filamentous structure (Figures 1&2) happened by a two-step process. In the first step, disproportionately high lipid synthesis compared to cytoplasmic volume increase (low growth rate during lag-phase) facilitated the transformation of the excess membrane into a bud (Figure 2D-F). Subsequently, these hollow buds detached from the inner membrane and were released into the periplasmic space (Figure 2E-F). In step two, these hollow vesicles fused and incorporated into the outer membrane, expanding into a hollow filamentous structure (Figures 1B & 2D-J).

The formation of these buds and associated membrane phase separation (Figure 4) can be explained by the model proposed by Lipowsky ^36^. It was hypothesized that when membranes phase-separate, more fluid L_d_-membrane spontaneously undergoes out-of-plane invagination and transforms into a bud to reduce line tension at the phase-boundary ^37^. The transformation of the L_d_-membrane into intracellular vesicles (Figure 4) observed in our study is in tune with this hypothesis. Nevertheless, apart from a more fluid L_d_-membrane, we also observed L_o_-membrane with high bending rigidity and low spontaneous curvature undergoing budding (Figure 4F). The formation of buds by the L_o_-membrane can be explained by the chemical composition of the cell membrane ^38^. In contrast to thermodynamic considerations, recent studies attempting to understand the behavior of cell membranes demonstrated the possibility of a more rigid L_o_-membrane undergoing budding. This selective invagination of L_o_-membranes was shown to have been induced by specific sterols within the membrane ^38^. In accordance with these studies, Gram-negative bacteria like *RS-P* and, in particular, anoxic phototrophs were known to have a diverse sterol composition in their plasma membrane ^39^, which could have played a role in the budding of L_o_-membranes.

Apart from the chemical composition, the nature of the membrane (L_o_ or L_d_) and the direction of its budding (into or out of the cytoplasm) could also have been determined by the differences in the macromolecular crowding between the cytoplasm and the periplasmic space ^22,40^. Previous studies attempting to understand budding and asymmetric cell division in aqueous two-phase LVs observed that phase separation of the aqueous core induced membrane phase separation and preferential association of L_d_ & L_o_-membrane with the aqueous phases of higher and lower densities, respectively. This preferential association between the membrane and aqueous phases was explained by the interaction between the membrane and the intracellular constituents ^41,42^. Such interactions were shown to alter the spontaneous curvature of the membrane and the direction of its budding ^41,42^. Our observation of membrane phase separation and subsequent budding in *RS-P* is in tune with these studies. L_d_-membrane in *RS-P* was always observed to have been in association with densely packed cytoplasm (∼17-35 wt% molecules), always budding into the cell and whereas L_o_-membrane preferentially invaginating out of the cell into less densely packed periplasmic space (∼6-7 wt% molecules) (Figure 4). These differences in the packing densities between the cytoplasm and the periplasmic space can also be inferred from the TEM images (Figure 2) ^43^.

Biophysical properties of the cell membrane and its phase separation into L_d_ & L_o_ phases was proposed to have played an essential role in the protocell reproduction ^40,44^. Nevertheless, such studies were primarily restricted to LVs (Figure 4). To our knowledge, ours is the first study to test these theories on self-replicating live cells. Unlike the studies that reported membrane phase separation in live cells^45^, we propose that the phase separation in *RS-P* was due to a passive reorganization of membrane components to reach a thermodynamically stable state rather than a functional membrane phase separation induced by an intracellular process or linked to a specific cellular function ^45,46^.

Morphologies of *RS-P* observed during log and stationary growth stages can also be explained from a physicochemical perspective. Growth (or expansion) of the cytoplasmic compartment within the cell, gradually filling the void left by the outer membrane, resembles LVs undergoing stretching when placed in an elongational flow path (Figures 2 & S3) ^29^.

Spherical LVs experiencing a low elongational strain were shown to transform into prolate or tubular shapes^47^ (Figures 3B, 3C & S3). However, at a certain strain level (threshold strain), vesicles transformed into a dumbbell shape. Upon further strain increase, vesicles formed elongated dumbbells with beaded structures appearing along their length. *RS-P* exhibited a similar pattern of morphological transformation (Figure 3 & S3). Like in LVs, morphologies of *RS-P* can be explained by the physical properties of the cell membrane, like its elasticity and expansion strain exerted on the membrane ^29,47^ due to cytoplasmic volume increase. In the case of *RS-P*, high and low elongational strains on the membrane translate into high and low growth rates (i.e., rate of cytoplasmic volume increase). In accordance with this hypothesis, short prolate or tubular cells were most frequently observed during the early lag phase, when the cell’s growth rate was low compared to other growth stages (Figure 3). With an increase in their growth rate during the mid/late-log phase, *RS-P* transformed into elongated dumbbells and, subsequently, dumbbell-shaped cells with beaded structures along their length (Figure 3).

The morphologies of *RS-P* observed in our study were similar to the LVs undergoing an osmotic deflation ^48,49^. When placed in a hyperosmotic solution, LVs undergo an osmotic deflation^50^ and transform into a plethora of morphologies, to attain a thermodynamically stable state. According to Area Differential Elasticity (ADE) theory ^48,49^, the end morphology of such vesicles is determined by the ratio of their surface area to volume. Although not undergoing osmotic deflation, *RS-P* is in a similar physical state due to excess membrane (Figure 2), i.e., high surface area: volume. However, in the case of *RS-P*, the surface area-to-volume ratio is everchanging and is determined by the growth stage (Figure 3). The morphologies of *RS-P* within a given growth stage and the change in its morphologies from lag, log, to stationary growth phases can be explained by the changing surface area-to-volume ratio. During the lag phase, the slow growth rate (rate of cytoplasmic volume) resulted in the formation of outer membrane extensions to accommodate the excess membrane.

During the early and mid-log phases, the cytoplasmic volume increase kept pace with the surface area increase of the cytoplasmic compartment, i.e., low surface area-to-volume ratio. This resulted in tubular cells with smooth surfaces or slight surface depressions (Figure 3A and S4). Cells with outer membrane extensions were rarely observed in the mid-log phase. Nevertheless, we do not presume this low ratio was due to the reduced rate of lipid synthesis. Closer observation of cells showed the presence of lipid droplets or hollow intracellular vesicles within the cytoplasm of the cells (Figure S4). Hence, we propose that the low surface area-to-volume ratio was due to the accommodation of excess lipids within the cytoplasm and the slow transfer of these lipids to the outer cell membrane. The extensive formation of intracellular vesicles also contributed to the rapid cytoplasmic volume increase in log-phase cells. A gradual decrease in the growth rate by the end of the log phase reverted cells to a state of excess outer membrane (Figure 1H&I). This transfer of lipids to the outer membrane transformed the cells from filamentous into a “string-of-beads” morphology.

A similar accumulation of lipids within the cytoplasm was observed when cells were kept in a constant state of logarithmic growth (Figure S5). Cells kept in a prolonged log phase gradually increased in length and diameter (Figure S5) but did not transform into “string-of-beads” morphologies. Such a transformation in these cells was only observed when their growth rate was reduced by stopping the transfers of these cells into fresh media. These results suggest that the growth rate had an influence on the morphology of the cells, and a faster rate of cytoplasmic volume increase reduced the rate of lipid transfer from the cytoplasm to the outer membrane. In spite of these observations, precise measurement of cell volume or the amount of excess membrane is technically challenging due to the lack of uniformity in cell morphology (Figure S10-S12). Moreover, from the mid-log phase, the cytoplasm of most cells had numerous intracellular vesicles (Figure S12). No uniformity was observed in the size, number, and morphology of these vesicles (Figures S4, S5, and S10-12).

Cell morphologies within any given growth stage were seldom uniform and, in general, were a mix of morphologies (Figure S10-S12). The length and width of the cells differed considerably from one incubation to another, even when cultivated under the exact same media composition. These morphological variations could be explained by the physiological differences among the cells within a population. Bacterial growth curves and their partition into different stages (lag, log & stationary), in general, represent the growth dynamics of the bacterial population. Nevertheless, averaging the data obscure considerable variation in the behavior of individual cells ^51,52^. It is known that genetically identical cells within a single bacterial population could exhibit considerable cell-to-cell variation in gene expression ^53,54^ and growth rates ^55^. The reason for such stochastic behavior of monoclonal cells has to date, not been well understood. Manifestation of these variations in the morphology of normal bacteria was not observed due to the presence of a rigid cell wall. Given the absence of a cell wall in *RS-P*, we assume such cell-to-cell variations in growth rate were manifested in its morphology (Figure S10-S12).

The morphological unpredictability of *RS-P* could also be explained from a bio-physical perspective. Recent studies attempting to understand similar unpredictability in LVs suggested that the initial aspect ratio (width: height) of the vesicle determines its end morphology^56,57^. An ever-changing aspect ratio of the vesicles resulted in unpredictable end morphologies. During the initial growth stages, we observed *RS-P* in a constant state of oscillation (ever-changing aspect ratio) (Video S1). We presume these oscillations could have originated within the cell due to asymmetric conditions on either side of the membrane ^21^ or random movement of cytoplasmic constituents (Video S2), or due to the presence of an internal metabolism ^58,59^. It was also hypothesized that internal metabolism could lead to shape instabilities on the cell surface ultimately leading to cell division ^58^.

A gradual reduction in the salt concentration of the media while maintaining a similar osmolarity with sucrose led to the transformation of *RS-P* back to its native form. This suggests that DSS played an essential role in stabilizing the protoplast state and preventing the formation of peptidoglycan cell walls ^60^. This also suggests that cells can reversibly transform from a stable protoplast state to their native state or *vice versa*, depending on the environmental conditions. Morphological differences between cells grown under different salt conditions (Figure 5) can be explained by the interaction between phospholipid membranes and cations ^24^. It was reported that divalent cations like Mg^+2^ organize lipids into clusters of 10-15 molecules each ^61^, dehydrating and stiffening the membrane and hindering its area expansion. On the other hand, monovalent cations like K^+^ reduce the bending rigidity of the membrane, making the membrane more flexible ^21^, conducive to the formation of larger cells. The absence of filamentous cells when *RS-P* was grown on a shaker (Figure 5I-L) was likely due to the constant shearing of long filamentous cells. As changes in cell morphology were reversible, when we cells were transferred between growth media containing different salt compositions, suggests that cell morphology was determined by the environmental conditions rather than the information encoded in its genome. In tune with the earlier studies that canonical structural proteins like FtsZ plays no role in regulating protoplast morphology ^14^, we did not observe an influence of FtsZ inhibitors on the lifecycles of *RS-P* (Figure S8).

During the late log phase, most *RS-P* cells are individual daughter cells formed by the fission of long filamentous cells. In the case of minimal cells or LVs, the proxy minimal cells, some earlier studies have hypothesized that the physiochemical forces of the environment could mediate such fission. Hence to achieve division, these studies employed methods like extrusion of LVs through a narrow pore size filter or application of an external force ^62–64^, to simulate the physical forces faced by such cells in the natural environments. Given that all our experiments are conducted under static conditions, we do not presume such forces could have contributed to the reproduction in *RS-P.* We instead propose that the force required to mediate fission was intrinsic to the cell. Recent studies showed that an internal metabolism or Brownian movement of intracellular constituents ^65,66^ could result in surface instabilities leading to cell division ^58,66^. In support of this presumption, cells in our incubations were intrinsically unstable (Video S1, early growth phase cells with expanded outer membrane).

They were in a constant state of oscillation, which should have resulted from the Brownian movement of intracellular constituents ^65^ (Video S1 & movement of DNA in spherical *RS-P* cells in the top half of Video S2). This constant movement of intracellular constituents could have led to the generation of force required for the fission of daughter cells in *RS-P* ^66^. In support of this presumption, we observed that cells within a filament were in a constant pull-and-push motion, which could have resulted in cell fission (Video S3 and Figure 1H-J).

The absence of intracellular constituents like DNA in the growth media suggests that loss of molecular biological coordination within the cells did not considerably affect the reproductive efficiency (Figure 6). Nevertheless, we do not presume *RS-P’s* reproduction was efficient. A considerable amount of membrane debris (Figures S6) and large clumps of cells in the stationary growth phase cultures (Figure 6C) suggest a lower reproductive efficiency than the cells in their original cell state. These clumps could have formed by the entanglement of filamentous cells (Figure S13). Observation of these clumps at regular intervals after their formation showed no evidence of them forming individual daughter cells and, over time, transforming into cell debris (Figure S13). Despite the wastefulness of *RS-P*’s reproduction in our static incubations, most daughter cells received DNA from the parent cell. They grew into filamentous cells upon transfer to fresh media, suggesting that this method of reproduction always resulted in metabolically viable daughter cells (Figure 6).

Given the absence of long filamentous cells and large cell clumps when grown on an orbital shaker (Figure 5I-L), we presume *RS-P* reproduced more efficiently under more turbulent environmental conditions (Figure 6E). Cells in natural environments are more likely to experience a similar turbulent force than being in a static state. The growth of *RS-P* daughter cells in media with glucose as the only carbon and energy source suggests that daughter cells received a full complement of genes from the parent cell to maintain an autonomous growth, capable of synthesizing all the required cell constituents from glucose alone (Figure S1).

Processes like growth and reproduction are punctiliously regulated by the timely expression of specific genes and precise localization of proteins within the cell ^67^. This thigh coordination between anabolic processes, growth, and reproduction is considered essential for the proper functioning of the cell ^68^. In contrast to this current understanding, we demonstrate that cells can still reproduce without such intracellular coordination. Excessive lipid synthesis, the multilamellar nature of the cell membrane, and favorable environmental conditions facilitated the formation of long filamentous cells by the sequential expansion of the outer membrane followed by the inner cell membrane. This two-step formation of long filamentous cells rather than large spherical cells by the random expansion of cytoplasm could have played an essential role in mediating leak-proof cell fission ^64^.

## Conclusions

In the present study, we demonstrated the influence of environmental conditions on protoplast reproduction. Our study shows that cells can reproduce independently of canonical molecular biological processes without significantly impacting reproductive efficiency under favorable environmental conditions. Given the simplicity and efficiency with which these cells reproduced, we propose primitive Gram-negative bacteria likely reproduced by a similar process before the evolution of complex intracellular processes.

## Acknowledgments

We want to thank Petra Schwille for her support and scientific input throughout the work and in preparing the manuscript. Gabriella Berthal and Markus Oster for excellent technical support. We thank the Advanced Light Microscopy Facility at EMBL, Heidelberg, Ulf Schwartz from Leica Microsystems, and colleagues at the departments of Ecological Microbiology (Bayreuth University) and of Cellular and Molecular Biophysics (Max Planck Institute for Biochemistry) for their support throughout the work.

## Author contributions

Conceptualization: DK, ML, BZ & TL. Culture experiments: DK. Electron microscopy: DK & AK. Light microscopy: DK & ML. Analysis: DK & JHK. Manuscript Writing: DK & JHK.

## Data Availability

Dr. Dheeraj Kanaparthi will share all data, materials, and methods upon reasonable request.

## Methods and Materials

### Cultivation of bacteria and generation of bacterial protoplasts

Gram-negative *Rhodobacter sphaeroides* (*DSM159*) was obtained from DMSZ (Braunschweig, Germany). Lyophilized cells were revived in nutrient broth (Sigma Aldrich, Germany) according to the procedure described by DSMZ. Protoplasts were generated from *R. sphaeroides* (*RS-P*) by inoculating overnight cultures into 3X nutrient broth (NB) containing 2% w/v sucrose, 80 µg/ml lysozyme, and 200µg/ml penicillin G (Sigma Aldrich, Germany). Cells were observed under a microscope to confirm the transformation of cells to their protoplast state. Over the next 3-4 weeks, the cells were repeatedly subcultured into fresh media with penicillin G and lysozyme, as described above. Subsequently, *RS-P* was transferred into NB with 7% Dead Sea Salt (DSS) (Amazon.de). The stability of these protoplasts (*RS-P*) was determined after ten sub-cultures by transferring cells into media without penicillin G or lysozyme. As *RS-P* did not revert to its native state with the cell wall under these conditions, lysozyme, and penicillin G were not added to the growth media for subsequent experiments. Unless otherwise stated, cells in the rest of the experiments were grown in 7%DSS-NB. In some experiments, we replaced DSS in the growth media with equivalent concentrations of other salts like MgCl_2_ and KCl (7%w/v). Cell growth was quantified at intervals by measuring their optical density at regular intervals (as mentioned in Figure 3) using a Biophotometer^®^ D30 (Eppendorf, Germany). The morphology of cells at different growth stages was determined by a variety of microscopic techniques (as described in the below sections).

### Influence of salt on the stability of protoplasts

The influence of salt on the stability of protoplasts was tested by two different methods. In the first method, the osmolarity of the media was reduced by dilution with NB. This was done in a step-by-step manner of replacing 10ml of the 7%DSS-NB (from the original 100ml) with 10ml of NB with no additional salt, day over seven days. All test and control experiments were replicated (n=6 each). To the control bottles, we added 10ml of the fresh 7%DSS-NB media. The morphology of the cells was observed using a phase-contrast microscope. No lysozyme and penicillin G were added to both of these media.

In the second method, we transferred 2ml of mid-log phase cells grown in 7%DSS-NB into 6xNB with 2% sucrose, 6xNB with 2% sucrose containing lysozyme and penicillin G and into 7%DSS-NB. 6xNB with 2% sucrose was used in these experiments to maintain the osmolarity similar to the media similar to 7%DSS-NB. All experiments were replicated (n=6 each).

To understand if the intracellular processes played a role in regulating the protoplast morphology, we inoculated *RS-P* into 7%DSS-NB with and without FtsZ functional inhibitor, 5mM 3-methoxybenzamide^30^. Cells were observed under a phase-contrast microscope at regular intervals for differences in the morphologies between the control and test incubations.

### Quantification of Superoxide dismutase activity and metabolic end products under aerobic growth

Minimal salt media (MSM) with 2% w/v glucose was used for these experiments. The composition of the minimal salt media was previously described by Kanaparthi et al., 2015 ^31^, with the exception of using 0.1g/l of yeast extract. 100ml of media was dispensed into 500ml Schott bottles (Schott AG, Germany). One set of growth media was made anaerobic using a vacuum manifold by repeatedly flushing the headspace with N_2_ gas and sealed with butyl rubber stoppers before autoclaving. After autoclaving, glucose and vitamin solutions were added to both aerobic and anaerobic media from the sterile anoxic stock solution. Aerobic and anaerobic media were inoculated (500μl) with overnight cultures of *RS*, grown in aerobic and anaerobic half-strength NB. A similar media composition was used to cultivate *RS-P* under aerobic and anaerobic conditions, except for adding 7%DSS. All the bottles were incubated on an orbital shaker at 30^0^C. To ensure uniform oxygen solubility, cultures grown under aerobic conditions were subject to shaking speeds of 150 rpm. All culture bottles were incubated in the dark to ensure non-phototrophic growth. All experiments were done in replicates (n=5). OD was determined every 3h interval for 15h. 1ml of the culture was centrifuged, and the supernatant was filter sterilized and stored at −20^0^C, for volatile fatty acid analysis. Glucose was quantified using a Glucose colorimetric assay kit (Invitrogen, Germany), according to manufacturers’ instructions.

SOD activity was determined using the 19160 SOD detection kit (Sigma Aldrich, Germany). Anaerobic culture bottles were transferred to an anaerobic chamber (Braun, Germany), and 1 ml of the culture was transferred into a 2 ml screw cap tube. The optical density of the cell was adjusted to a similar value in all the samples. Cells were homogenized using silica beads (Sigma Aldrich, Germany) and vertexing for 3min. All tubes were briefly centrifuged, and 250μl of the cell lysate was transferred into the microtiter plate, and other reagents were added according to the manufacturer’s instructions. The plates were then transferred out of the anaerobic chamber and incubated at 37^0^C for 1h. The colorimetric measurements were done using a 96-well plate reader (Thermo Fischer Scientific, Germany). A similar experimental procedure was used for determining the SOD activity of *RS* and *RS-P* cells grown under aerobic conditions. All the experiments were done in replicates n=10 for the data presented in the manuscript, but the experiment was repeated several times over the course of our work.

### Phase-contrast and Super-resolution STED microscopy

Morphology *RS-P* was determined at regular intervals using an Axioskop 2plus microscope (Carl Zeiss, Germany) using a Plan-NEOFLUAR 100X/1.3 objective. Images were acquired with Leica DSF9000 (Leica, Germany) cameras. STED microscopy was performed with an inverted TCS SP8 STED 3X microscope (Leica Microsystems, Mannheim Germany) using an 86x/1.2 NA water immersion objective (Leica HC PL APO CS2 - STED White).

Fluorophores were excited with either 488, 561nm, 594nm, or 633m laser light derived from an 80 MHz pulsed White Light Laser (Leica Microsystems, Mannheim Germany). For stimulated emission, either a pulsed 775 nm laser or a 592nm CW laser (Leica Microsystems, Mannheim, Germany) was used depending on the fluorophore. The emitted fluorescence light was collected with Hybrid Detectors (HyD, Leica Microsystems, Mannheim Germany) using appropriate emission band pass-filter setting and typically a gate of 0.3ns-6.0ns for depletion with the 775nm laser or 1.0-6.0ns with the 592nm laser. Images were recorded in photon counting mode and line accumulation. Image deconvolution was performed on selected images and video with Huygens Professional (version 16.10.1p2, Scientific Volume Imaging, Hilversum, The Netherlands).

### Transmission electron microscopy

*RS-P* cells at different growth stages were harvested from the growth media by centrifugation at 500rpm for 5min. Cells were then preserved in 2.5% v/v glutaraldehyde (Carl Roth, Karlsruhe, Germany). Cells were subsequently stained with 0.25% w/v uranyl acetate. Cells were subsequently dehydrated in increasing concentrations of acetone. Cells were embedded in epoxy resin and left for polymerization for 72h. Epoxy blocks were sectioned, and cells were counter-stained with 1% lead citrate. TEM was carried out using a Zeiss EM 912 (Zeiss, Oberkochen) with an integrated OMEGA filter at 80kV. Images were acquired with 2k x 2k pixel slow scan CCD camera (TRS, TrÖndle Restlichtverstrkersysteme, Moorenweis, Germany) and an ImageSP software (version 1.2.9.77) (SysProg, Minsk, Belarus).

### Quantifying reproductive efficiency and metabolic viability of daughter cells

Reproductive efficiency was determining the leakage of cell constituents during the formation of daughter cells. The viability of the daughter cell was determined by staining and quantifying cells by flow cytometry.

*RS-P* was inoculated into nutrient broth containing 7% w/v of DSS, NaCl, or MgCl_2_. The culture bottles were incubated under static conditions or on an orbital shaker at 160 rpm. 2ml of cells were harvested every 24h intervals from all the incubations. In the case of static cultures, incubations were briefly placed on an orbital shaker for 10 min at 100 rpm to disperse the biofilm. Intact cells were precipitated by centrifugation at 6000 rpm for 10 min. DNA was extracted from both the cell and supernatant. In the final step, DNA was washed with 70% ethanol three times to remove any residual salts from DNA. The resultant DNA was dissolved in 2ml of TE buffer. DNA was quantified using the Qubit dsDNA quantification kit (Invitrogen). All the experiments were done in replicates (biological replication, n=5).

The viability of daughter cells was determined by the growth of daughter cells either by directly sub-culturing 10-20 day old cultures into fresh media or passing 10-20 day old cultures through 0.45μm cellulose acetate filter (Millipore) and then transferring the filtrate into new media. The presence of intracellular DNA and cytoplasmic activity of daughter cells was also tested by staining them with DNA stain, PicoGreen^TM^ (Invitrogen), and live cell stain CellTrace^TM^ violet stain (Invitrogen). Cells with intracellular DNA and cytoplasmic activity were quantified using flow cytometry (Attune NxT, Thermo Fisher Scientific).

