## Supplement for "The reproduction of Gram-negative protoplasts and the influence of environmental conditions on this process"

### Supplementary Materials

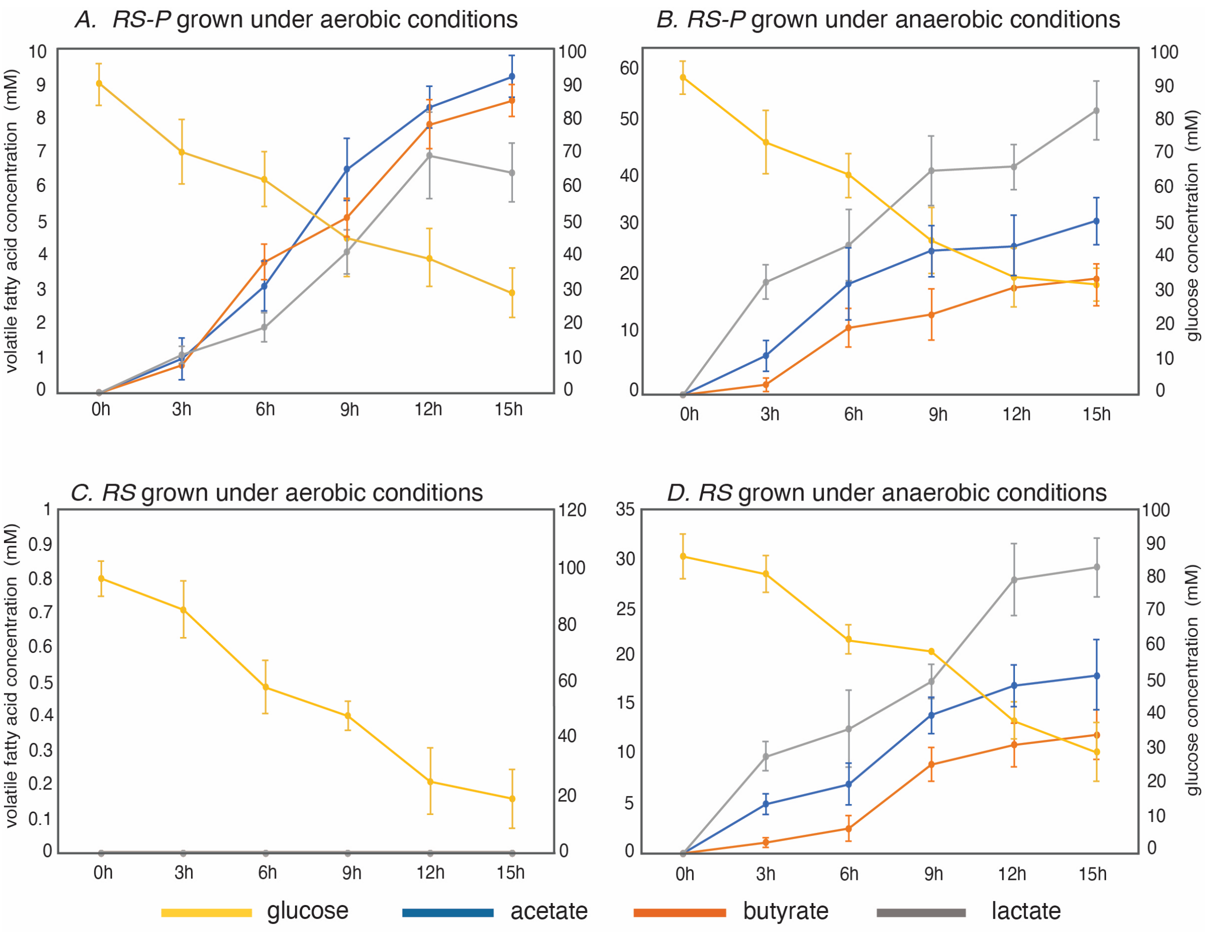


Figure S1. Glucose consumption by *RS-P* and *RS* under aerobic and anaerobic conditions, related to Figure 3.

The plot shows a gradual consumption of glucose and an increase in volatile fatty acids like lactate, butyrate, and acetate during the growth of *RS-P* and *RS*. The X-axis shows time (hours), the primary Y-axis shows the concentration of volatile fatty acids (acetate, lactate, and butyrate), and the secondary Y-axis shows the concentrations of glucose. All the measurements were cone in replicates (n=5).

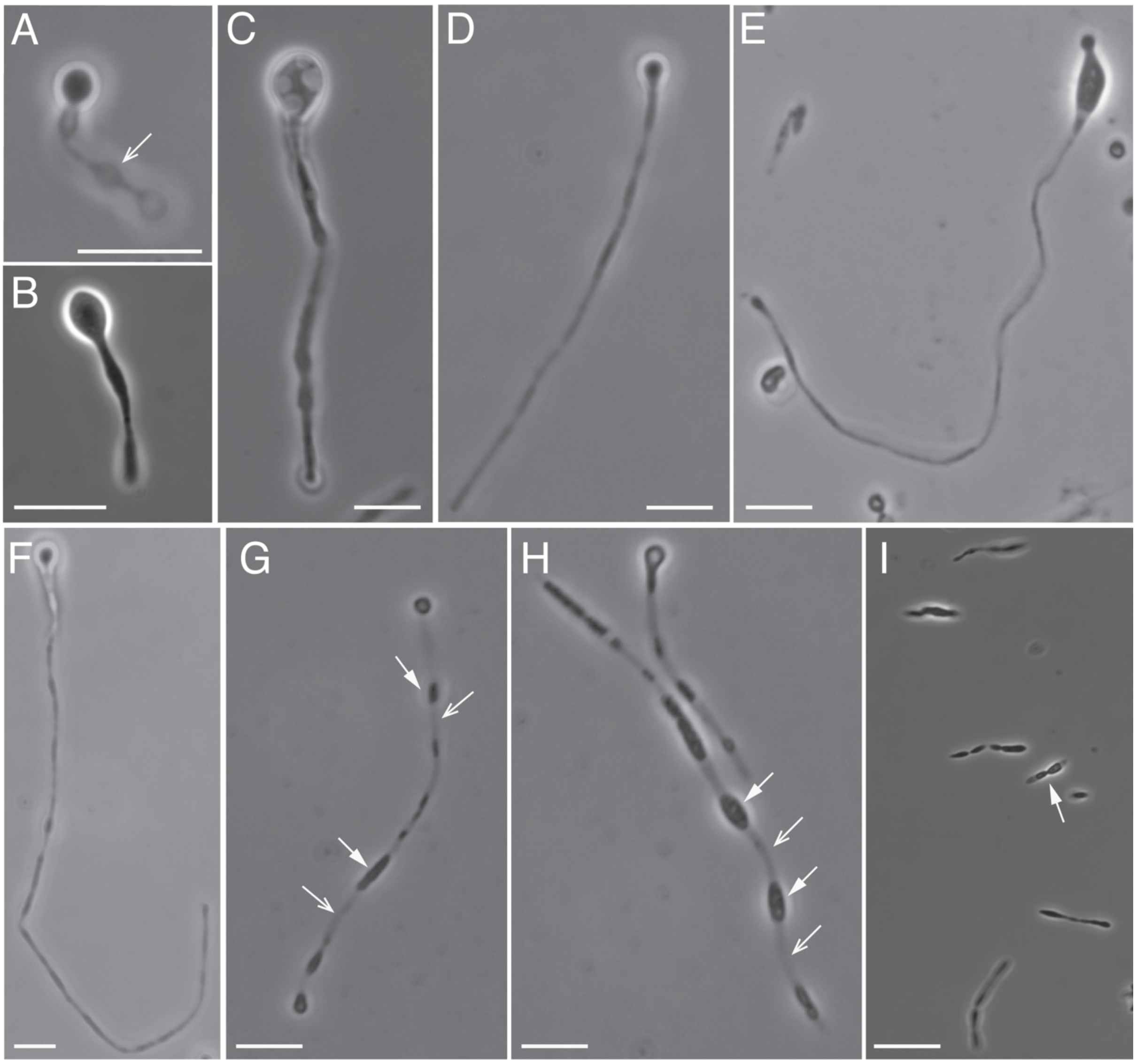


Figure S2. The life cycle of *RS-P*, related to Figure 1.

Images A – I show phase-contrast images of *RS-P* during different growth stages. The arrow in image A points to the hollow outer membrane extension. Image B – F shows subsequent growth stage filamentous cells. Images G & H show stationary growth phase cells with the restricted presence of cytoplasm. Closed and open arrows in these images point to regions of the filament with and devoid of cytoplasm. The image I show cells in their last growth stage. Arrows in this image point to cells formed by the fragmentation of long filamentous cells. Scale bars: 5µm.

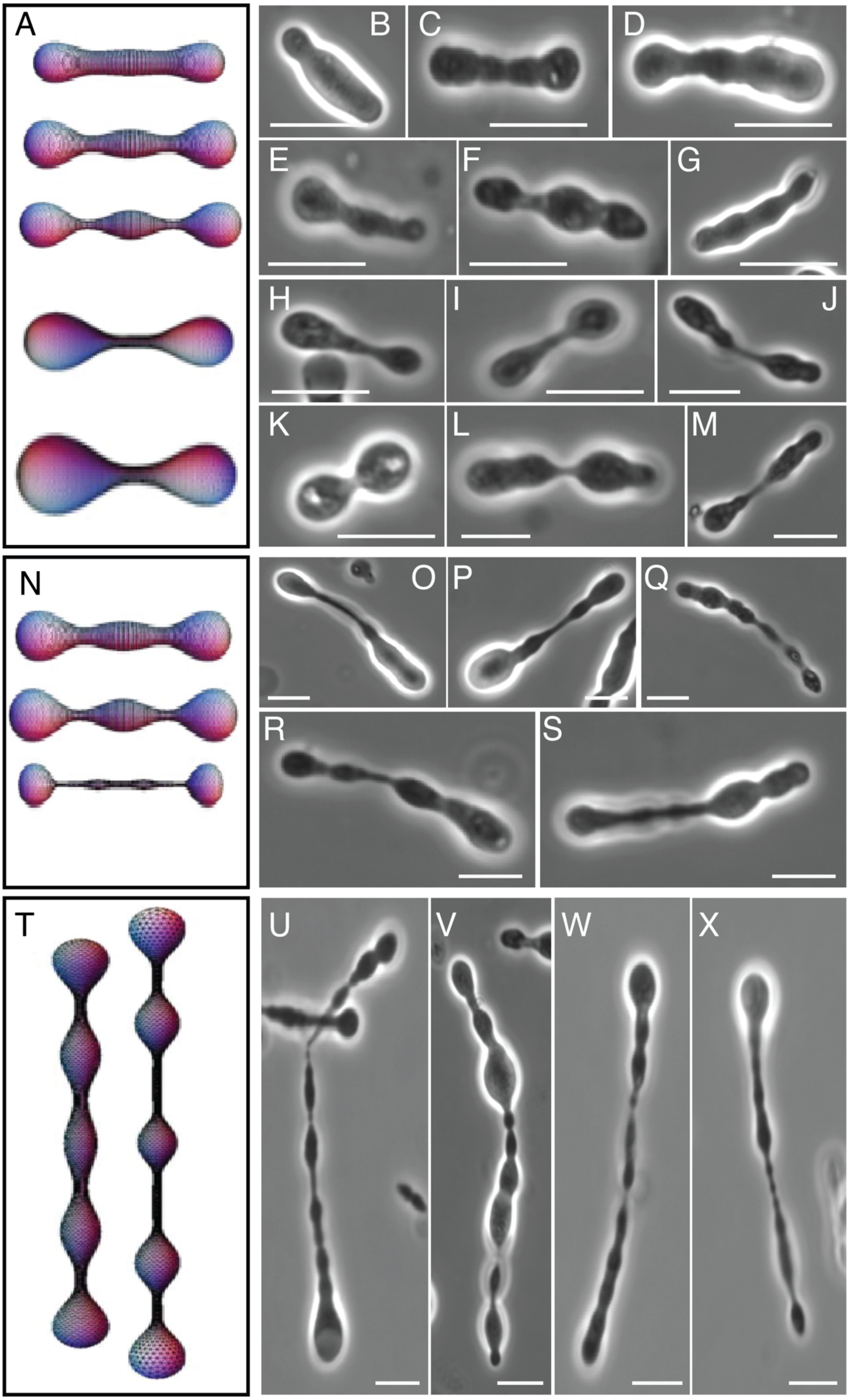
A
B
C
D
E
F
G
H
I
J
K
L
M
N
O
P
Q
R
S
T
U
V
W
X

Figure S3. Comparison of lipid vesicle morphologies with *RS-P*, related to Figure 3.

Images A, N & T are theoretically predicted morphologies of vesicles undergoing stretching in the laminar flow path (originally published by Narasimhan et al., 2015, reproduced here with permission from Cambridge University Press) *(23)*. Image A: Vesicle subjected to low elongational stress. Images B – M are early growth phase *RS-P* cells. Image N shows vesicles subjected to higher elongational stress. Images O – S show log phase *RS-P* cells. Image T shows vesicles subjected to even higher elongational stress. Images U – X show late log and early stationary phase *RS-P* cells. Scale bar: 2μm.

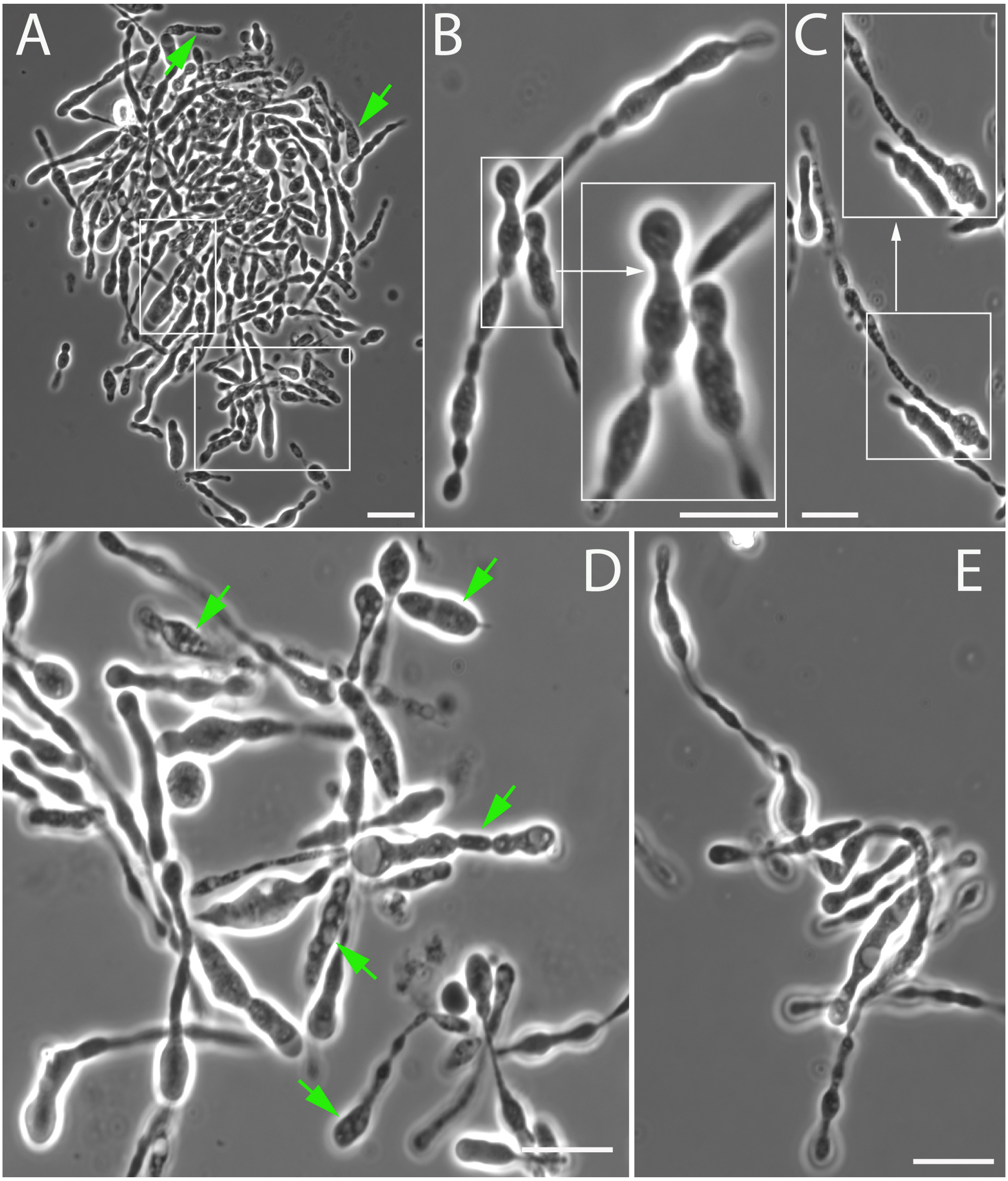


Figure S4. *RS-P* cells in the log phase, related to Figures 1 & 2.

Images A-E show log phase *RS-P* cells. Magnified regions in A-C and arrows point to intracellular vesicles in *RS-P*. These cells also exhibited slight surface deformations, suggesting excess membrane. Scale bar: 10μm.

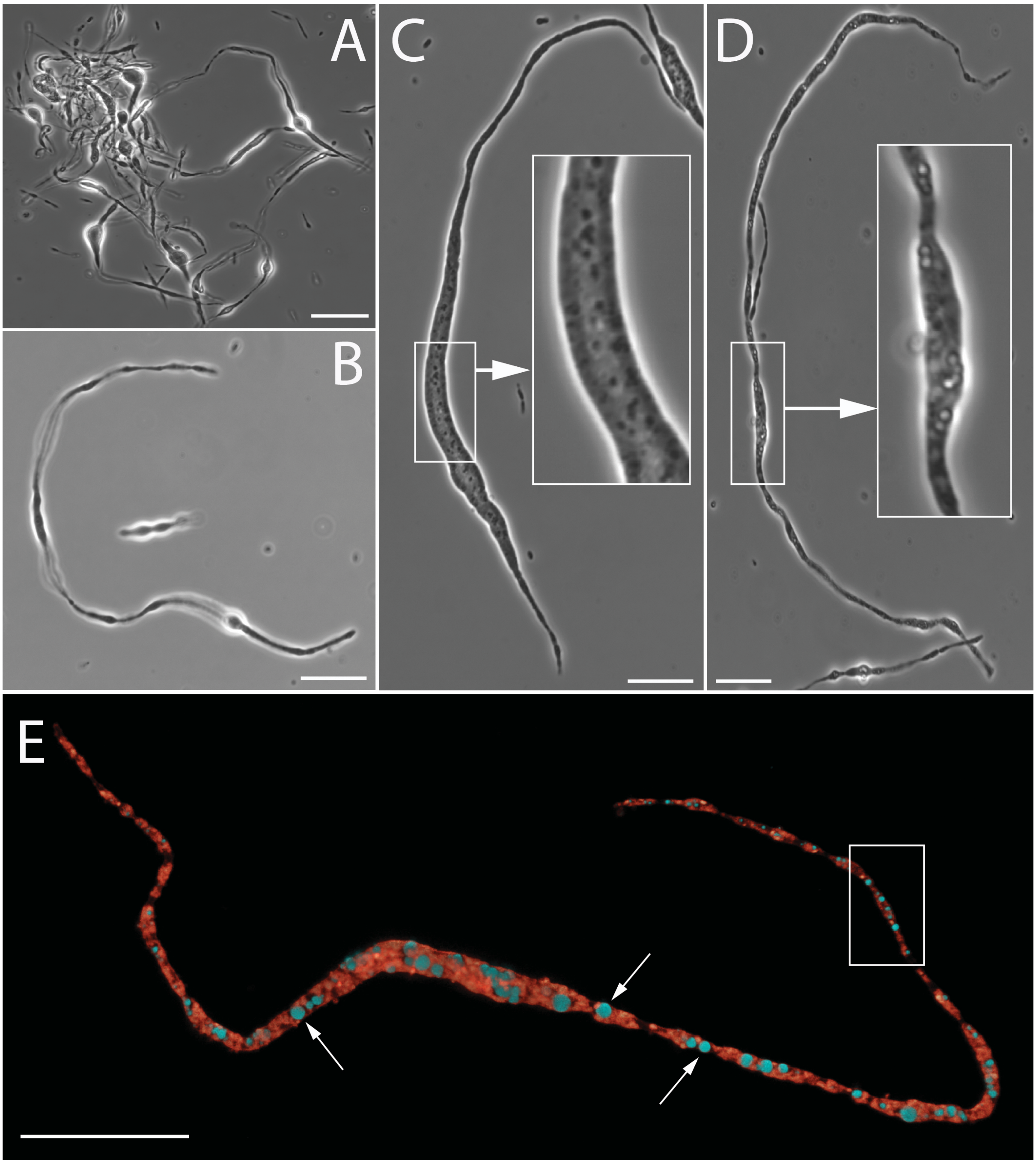


Figure S5. *RS-P* cells in a continuous log phase, related to Figures 1 & 2.

*RS-P* cells are sub-cultured every few hours to keep them in a constant state of logarithmic growth. Under these conditions, cells rarely developed surface depressions and transformed into a “string-of-beads” morphology. The highlighted regions in images C-E show excess membrane within these cells in the form of hollow intracellular vesicles. Image E is the

STED microscope image of the *RS-P* cell. Cells in these images were stained with FM™5-95 (all membrane, red) and *FAST*™ DiI ($L_d$-membrane, green). Scale bar: 10µm.

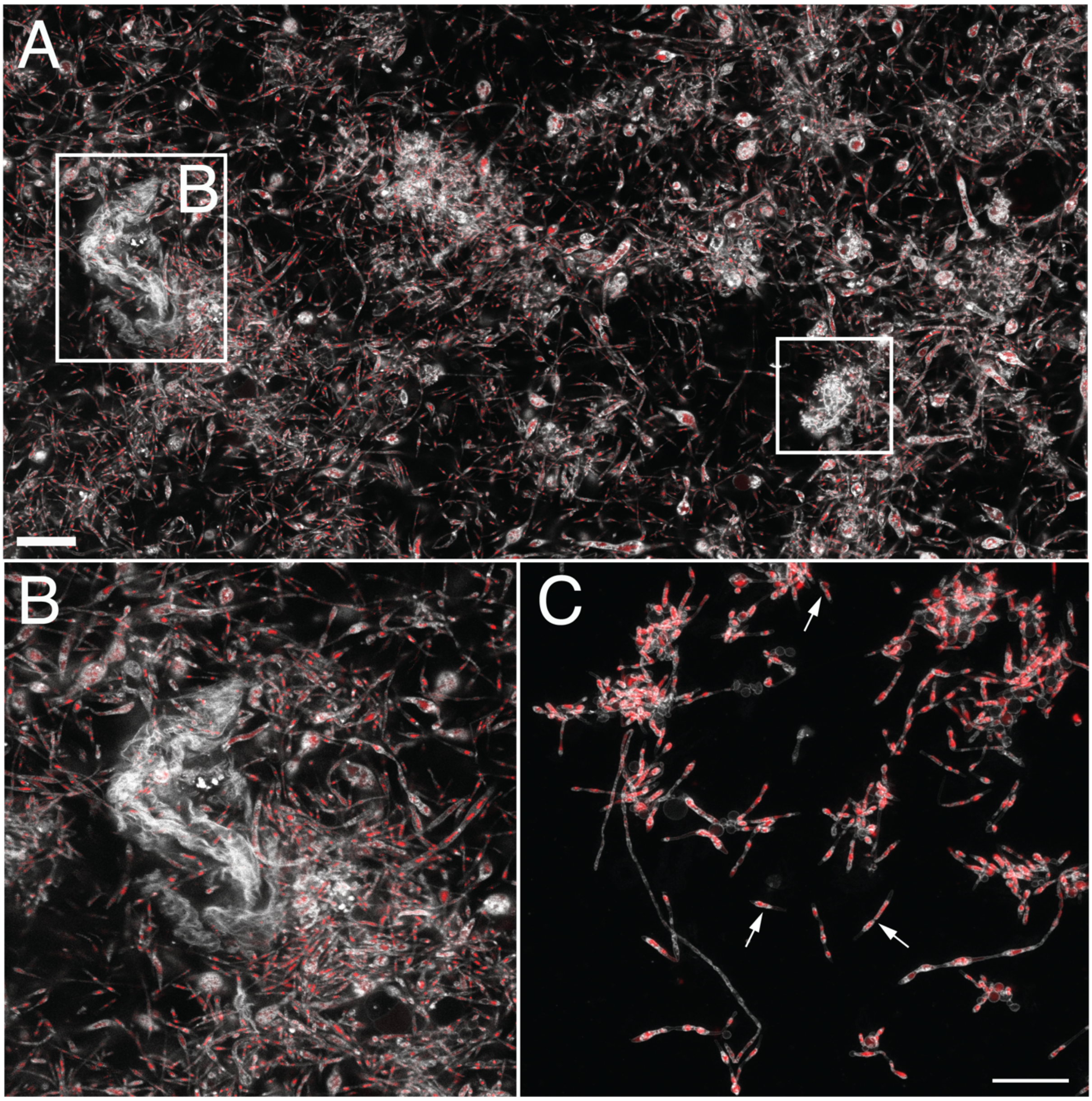


Figure S6. Stationary growth phase *RS-P* cells, related to Figures 1 & 2.

The image shows *RS-P* cells in the stationary growth phase. Cells were stained, imaged, and color-coded, as in Figure 1. Image A shows a wide-field image of early stationary growth phase *RS-P* cells. All cell morphologies, like tubular, filamentous, and individual daughter cells, can be seen in the image. Image B is the magnified region of membrane debris observed during the stationary growth phase. Image C is the close-up view of filamentous *RS-P* cells during the late stationary growth phase. Arrows in image C point to individual daughter cells or cells formed by the fragmentation of filamentous cells. Scale bar: 30μm A and 10μm C.

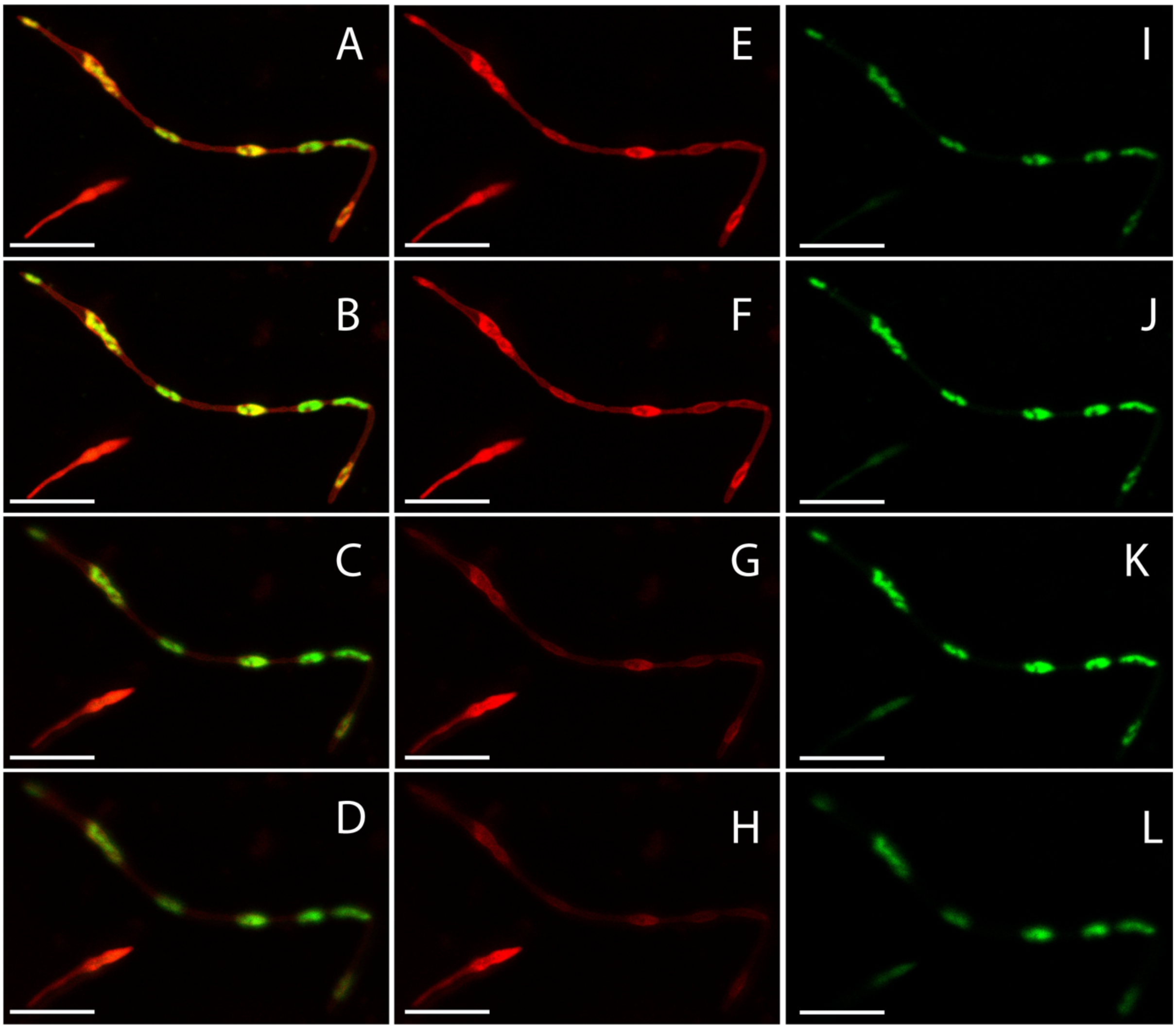


Figure S7. Membrane phase-separation in *RS-P, related to Figure 4.*

Cells in these images were stained with FM™5-95 (all membrane, red) and *FAST*™ Dil ($L_d$-membrane, green). Images A - D: Successive optical sections of the same *RS-P* cells (along the Z-axis, 0.2μm intervals). Images E – L show images of the same cell in individual channels. Scale bar: 10μm.

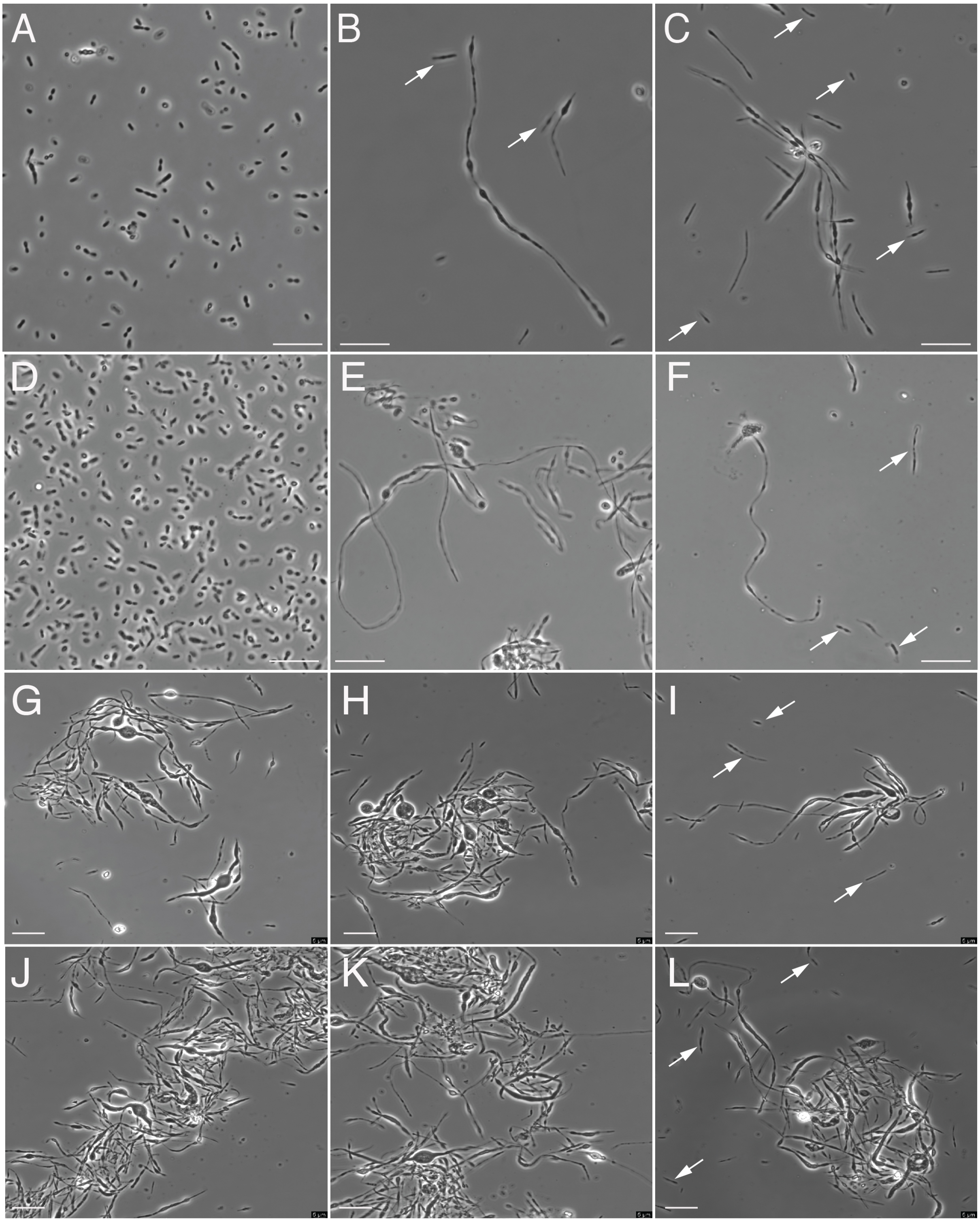


Figure S8. Influence of 3-methoxybenzamide on the morphology of *RS-P*, related to Figure 5.

Images A-L are phase contrast images of *RS-P* grown in media with and without the FtsZ activity inhibitor 3-methoxybenzamide. Images A-C are images of *RS-P* cells grown in 7%$MgCl_2$-NB. Image A and C&D show early, and mid-growth stage *RS-P* cells, respectively. Images D-F are images of *RS-P* cells grown in 7%$MgCl_2$-NB with 3-methoxybenzamide. Image D and E&F show early, and mid-growth stage *RS-P* cells, respectively. Images G-I are the images of *RS-P* cells grown in 7%DSS-NB. Image G and H&I show early, and mid-growth stage *RS-P* cells, respectively. Images J-L are images of *RS-P* cells grown in 7%DSS-NB with 3-methoxybenzamide. Image J and K&L show early, and mid-growth stage *RS-P* cells, respectively. Scale bars: 10μm.

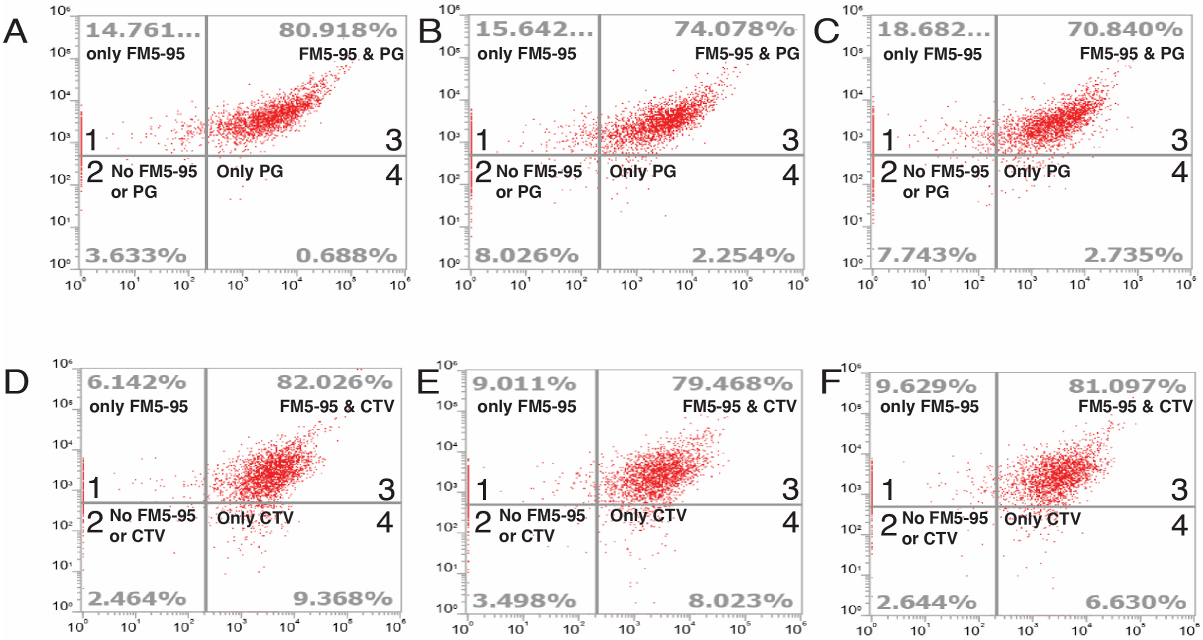


Figure S9. Quantification of DNA content and cell integrity of *RS-P* daughter cells by flow cytometry, related to Figure 6.

A-C: Staining and quantifying cells with an intact membrane and intracellular DNA (triplicates). Quadrants represent cells classified positive for only $FM^{TM}$5-95 (quadrant 1), negative for both dyes (quadrant 2), positive for both $FM^{TM}$5-95 and $PicoGreen^{TM}$ (quadrant

3), and positive only for PicoGreen™ (quadrant 4). These results show that most daughter cells (70-80%, quadrant 3) had intact membrane and received DNA from the parent cells. D – F: Staining and quantifying cells with an intact membrane and cytoplasmic activity (triplicates). Quadrants represent cells classified positive for only FM™5-95 (quadrant 1), negative for both FM™5-95 and CellTrace Violet (quadrant 2), positive for both FM™5-95 and CellTrace Violet (quadrant 3), and positive only for CellTrace Violet (quadrant 4). These results show that most daughter cells (79-82%, quadrant 3) had intact membrane and exhibited cytoplasmic enzyme activity.

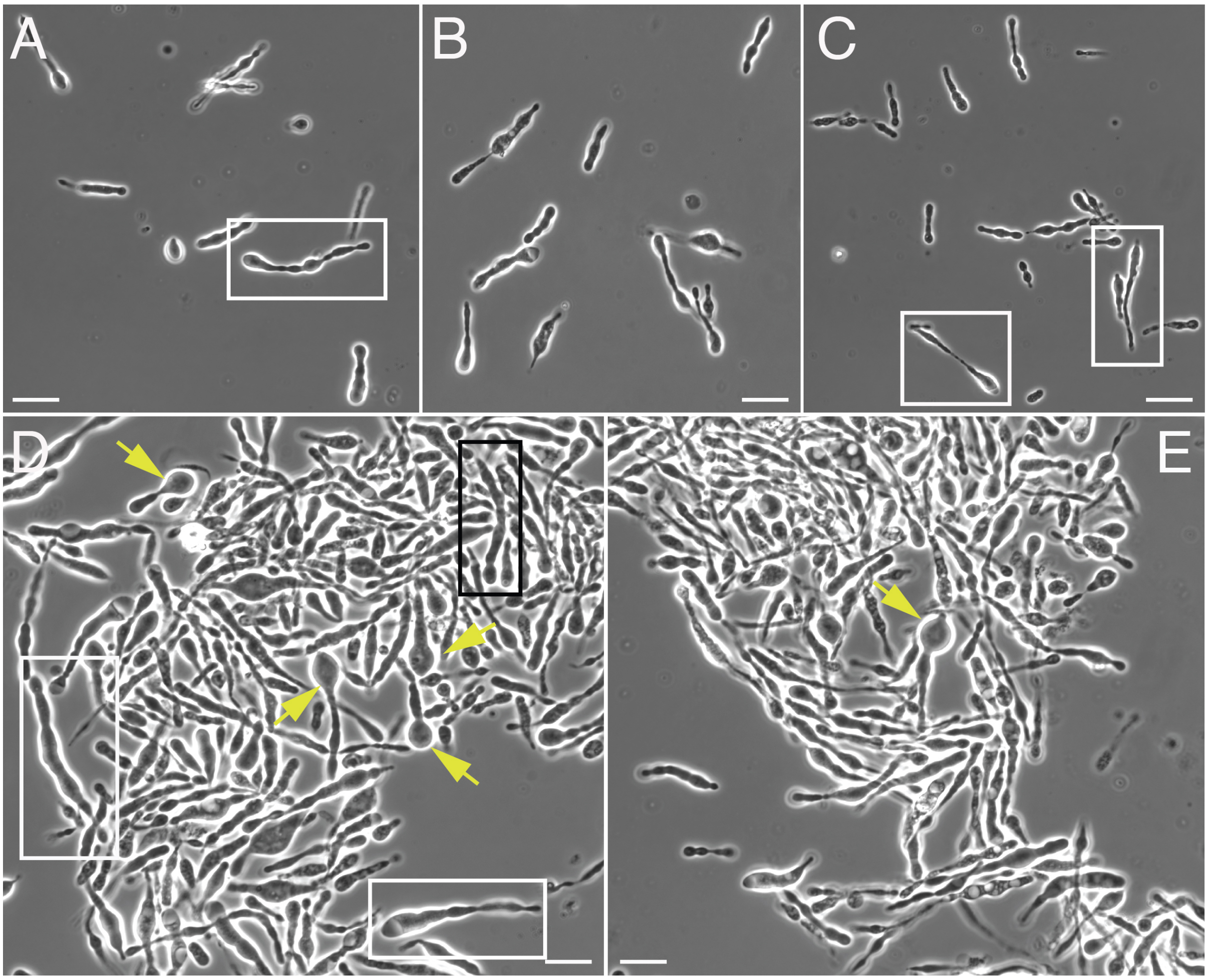


Figure S10. Morphological variation mid-log phase in *RS-P*, related to related to Figures 1 & 2.

Images A-E show images of *RS-P* from a single microscopic slide. Cells within the boxed regions (black) highlight variations in the diameter of the filamentous cells. Arrows point to the filamentous cells with lenticular structures. Scale bar: 10μm.

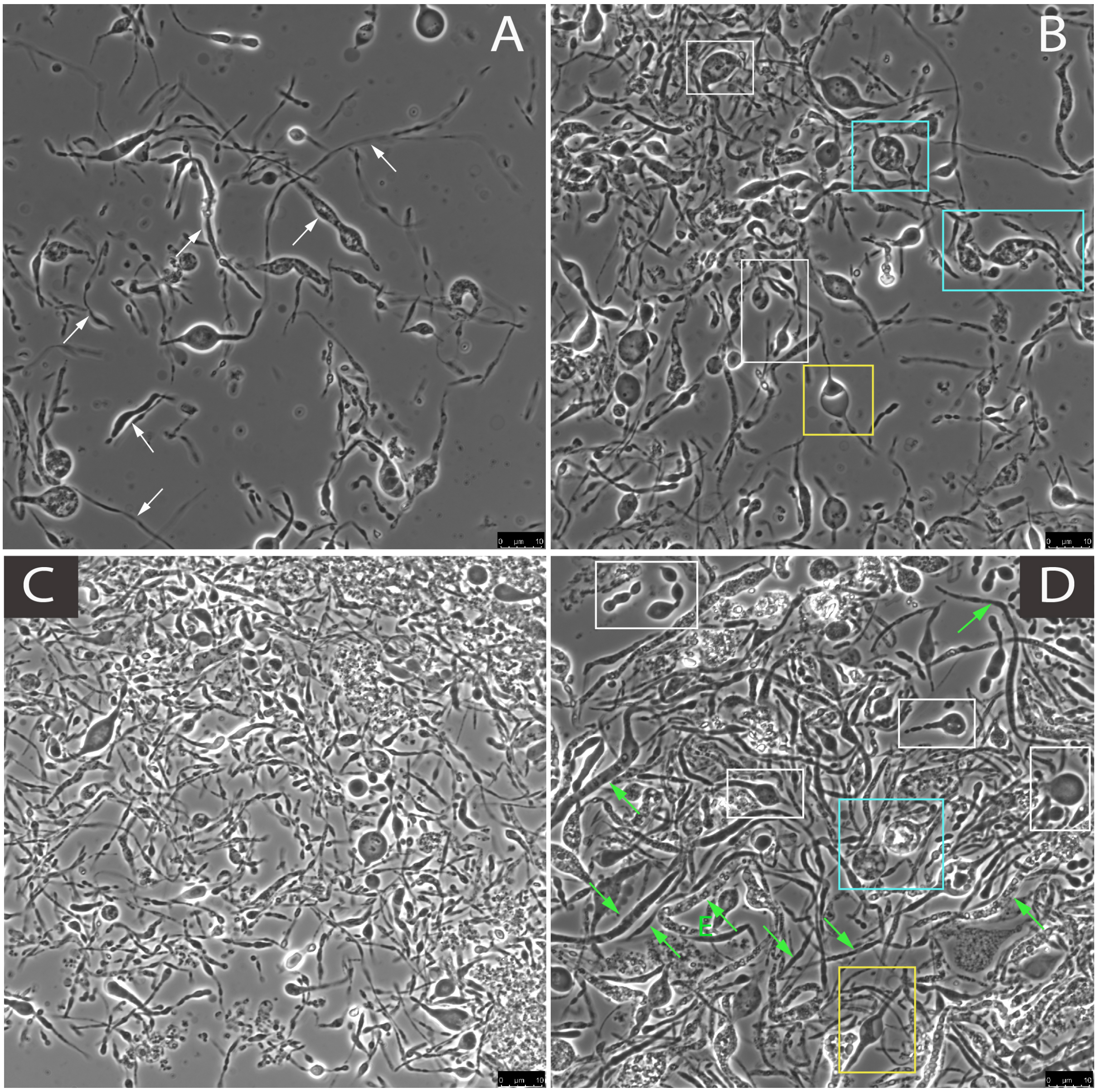


Figure S11. Morphological variation late log phase in *RS-P*, related to Figures 1 & 2. Images A-D show phase-contrast images of *RS-P*. Arrows in image A point to filamentous cells. Boxed regions in image B&D highlight the lenticular structures within the filaments. Lenticular structures within the yellow boxes exhibit a single large intracellular vesicle. Cells within the blue boxes exhibit multiple hollow intracellular vesicles. Green arrows in image D point to the filamentous cells that differ in diameter. A similar lack of morphological uniformity among cells can be seen in image C. TEM images of such cells with intracellular vesicles are shown in Figure S12. Scale bar: 10μm.

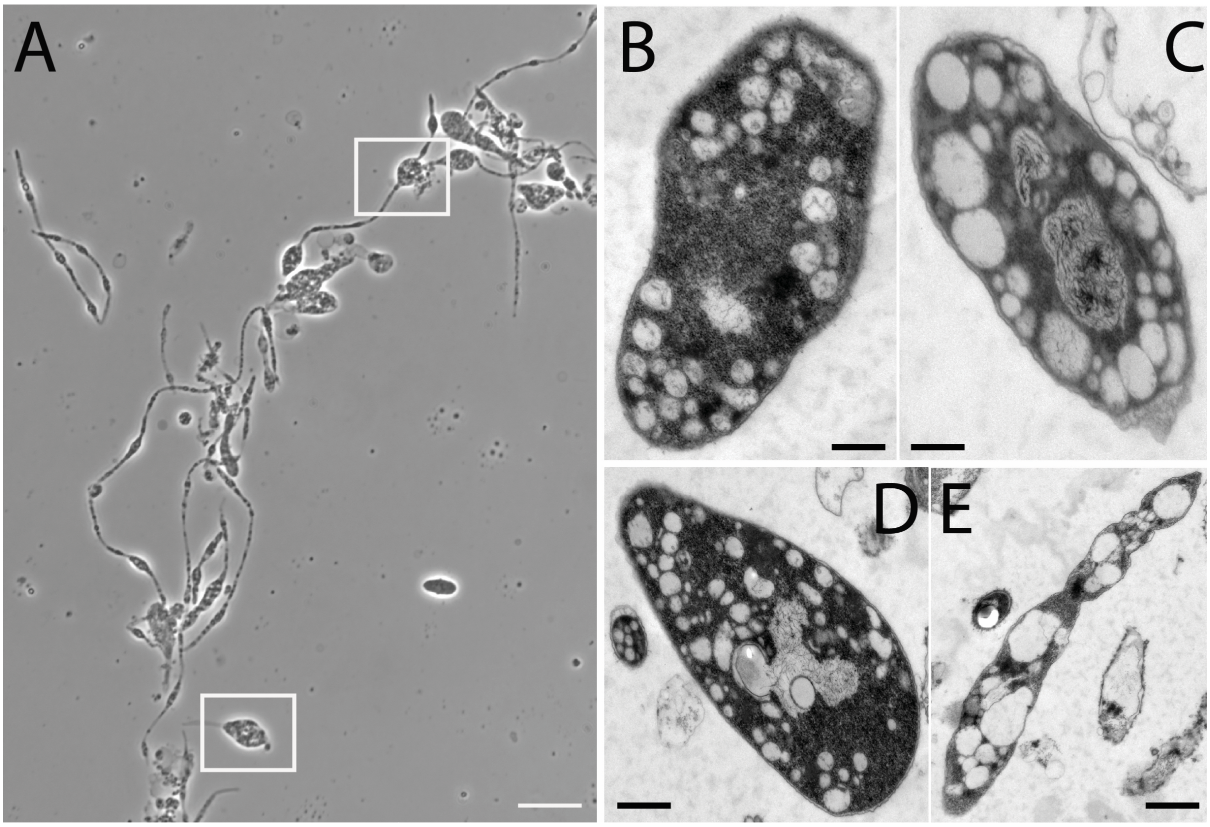


Figure S12. Filamentous *RS-P* cells with intracellular vesicles, related to Figures 3 & 4. Image A and images B-E show filamentous *RS-P* cells with intracellular vesicles. No uniformity was observed in the size or number of hollow vesicles within the cell. Scale bar: 10μm (A) & 250μm (B-E).

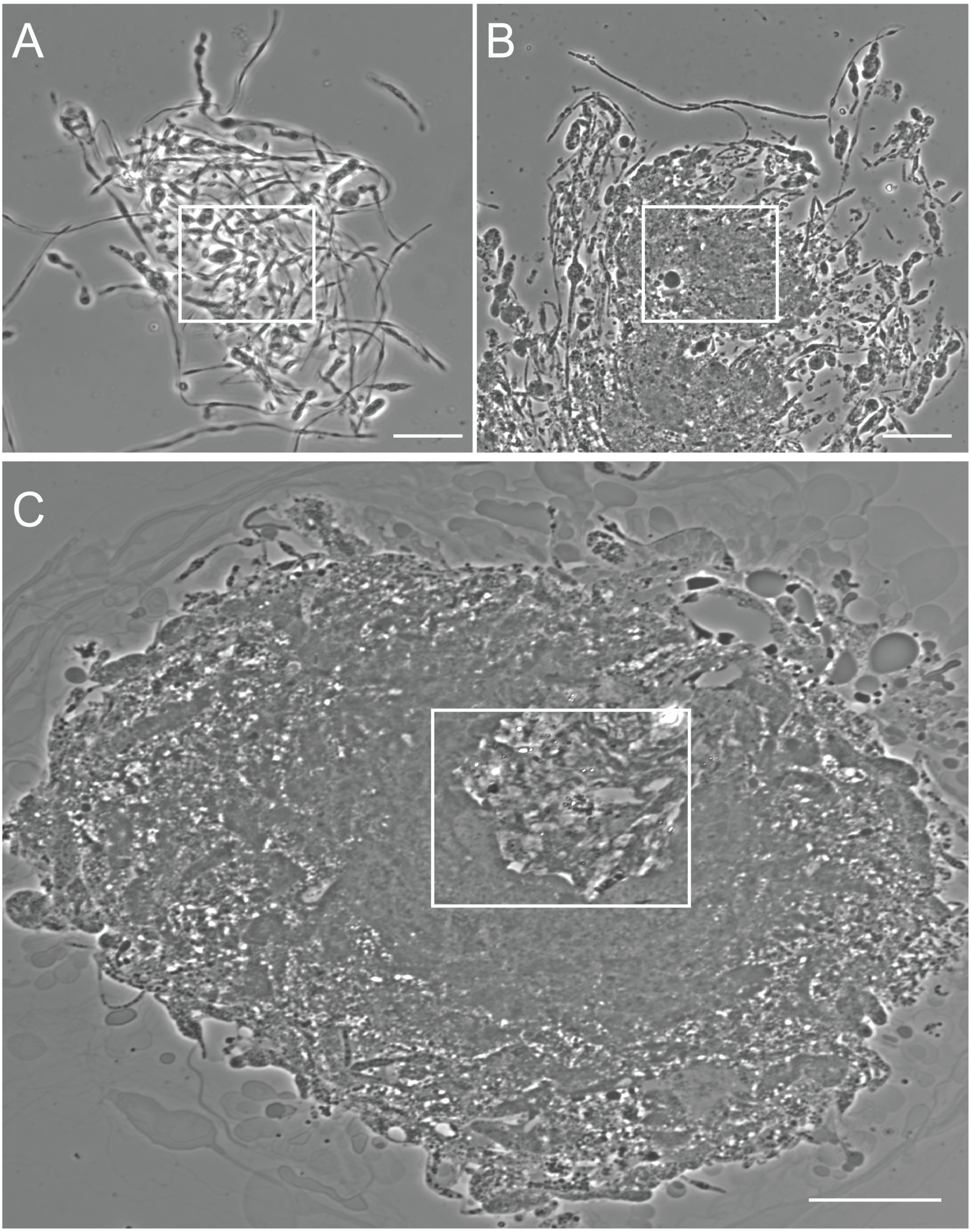


Figure S13. Sequential steps involved in the formation of cell clumps, related to Figure 6. Image A shows entangled filamentous cells during the log phase. Image B shows stationary growth phase cells. The highlighted region in the image shows an indistinguishable mass of the *RS-P* cell. Image C shows cells in the subsequent growth stage. The highlighted area in the image shows the central region of the clump where lipids from the degraded cells self-assembled into lipid layers. Lipids extruding through the periphery of the clump can also be seen in image C. Scale bars: 10μm.
